# Her6 and Prox1 are novel regulators of photoreceptor regeneration in the zebrafish retina

**DOI:** 10.1101/2023.03.20.532714

**Authors:** Kellie Veen, Aaron Krylov, Shuguang Yu, Jie He, Patrick Boyd, David R. Hyde, Theo Mantamadiotis, Louise Y Cheng, Patricia R Jusuf

## Abstract

Damage to light-sensing photoreceptors (PRs) occurs in highly prevalent retinal diseases. As humans cannot regenerate new PRs, these diseases often lead to irreversible blindness. Intriguingly, animals, such as the zebrafish, have the ability to regenerate PRs efficiently and restore functional vision. Upon injury, mature Müller glia (MG) undergo reprogramming to adopt a stem cell-like state. This process is similar to cellular dedifferentiation, and results in the generation of progenitor cells, which, in turn, proliferate and differentiate to replace lost retinal neurons. In this study, we tested whether factors involved in dedifferentiation of *Drosophila* CNS are implicated in the regenerative response in the zebrafish retina. We found that *hairy-related 6* (*her6*) negatively regulates of PR production by regulating the rate of cell divisions in the MG-derived progenitors. *prospero homeobox 1* (*prox1*) is expressed in differentiated PRs, and likely promotes PR differentiation through phase separation. Interestingly, upon Her6 downregulation, Prox1 is precociously upregulated in the PRs, to promote PR differentiation; conversely, loss of Prox1 also induces a downregulation of Her6. Together, we identified two novel candidates of PR regeneration that cross regulate each other, and may be exploited to promote human retinal regeneration and vision recovery.

## Introduction

Vision is initiated in the eye by specialised light-detecting cells termed photoreceptor (PR) neurons. PRs are sensory neurons that convert light into electrical signals that are then transmitted to subsequent neural circuits and interpreted by the brain (Noel et al., 2021). PRs can be classified as either rods, which detect low levels of light, allowing for low-level scotopic vision such as night vision; or cones, which detect specific wavelengths of light at greater intensities, conferring photopic vision such as colour vision. Rods and cones are highly specialised neurons consisting of complex and precise structures that each carry out specific functions. Both PR types are comprised of a light-sensitive outer segment, a connecting cilium, a cell body, and a synaptic terminal. Due to the complex nature of PR development, as well as their high metabolic and functional demands, PRs are vulnerable to mutations and prone to visual diseases. In the disease retinitis pigmentosa, PRs within the retina degenerate over time (Parmeggiani, 2011). As human retinal cells cannot regenerate, this vision loss is irreversible. Unlike humans, zebrafish retinae are able to regenerate after injury. Therefore, it is key to understand how zebrafish regeneration occurs, such that potential therapies to regenerate lost human PRs can be developed.

Zebrafish regeneration is achieved by the reprogramming of mature Müller glia (MG) into a stem-like state, and subsequent production of progenitor cells that differentiate into mature neurons such as PRs (Bernardos et al., 2007, D’Orazi et al., 2020, Fraser et al., 2013, Montgomery et al., 2010, Hagerman et al., 2016, Ng et al., 2013). Dedifferentiation, the process in which mature cell types are reprogrammed and adopt stem-like properties (Friedmann-Morvinski et al., 2012, Southall et al., 2014, Froldi et al., 2015), is a crucial mechanism in regeneration, as it enables the acquisition of a stem-like fate in mature cells. Therefore, dedifferentiation is a key step in the regeneration of PRs, and factors implicated in this process may be involved in regeneration.

In this study, we found two transcription factors implicated in dedifferentiation of the *Drosophila* brain to be involved in regeneration of the zebrafish retina. Hairy-related 6 (Her6), the zebrafish ortholog of *Drosophila* Deadpan (Dpn), and Prospero homeobox 1 (Prox1), the zebrafish ortholog of Prospero (Pros) mediate the regenerative response in zebrafish PRs. The regenerative response involves MG acquisition of stem cell fate, proliferation, migration, differentiation, and integration into the functional circuit (Lahne et al., 2020). We found that Her6 is a negative inhibitor of progenitor cell divisions, whereas Prox1 is required to promote PR differentiation via phase separation in the context of retina regeneration. Her6 is downregulated in the MG-derived progenitors and its knockdown was sufficient to increase the number of PRs after injury, suggesting that Her6 is a negative regulator of regeneration. Prox1 was expressed in the differentiating PRs and knockdown of Prox1 resulted in reduced PR differentiation, mediated through Liquid-Liquid Phase Separation (LLPS). Taking these observations together, we propose novel roles for Her6 and Prox1 in PR regeneration.

## Materials and methods

### Fly husbandry and strains

*Drosophila melanogaster* stocks were maintained on standard medium at room temperature (22°C). Overexpression and knockdown experiments were set up at 25°C, and after 48 hours, the progeny was heat shocked for 15 minutes at 38°C and moved to 29°C.

The fly strains used were: hsFLP, act>CD2> Gal4; UAS-Dcr2, UAS-GFP/ TM6b, prospero RNAi/ CyoYFP; UAS GFP/ MKRS, UAS-deadpan (A. Baonza), UAS-elav RNAi (#37915, VCRC), UAS-mCherry RNAi (#35785, Bloomington), UAS-SoxNeuro RNAi (#25996, Bloomington).

### Immunostaining fly samples

At five days after larval hatching, larvae were dissected in phosphate buffered saline (PBS), fixed in 4% formaldehyde for 20 minutes, and washed in PBS containing 0.5% Triton X 100 (PBST). Tissues were incubated in a primary antibody solution overnight at 4°C, followed by an overnight incubation of secondary at 4°C. Samples were mounted in 80% glycerol in PBS for image acquisition. Primary antibodies used were mouse anti-Mira (1:50; gift of Alex Gould) and chick anti-GFP (1:2000; Abcam). Secondary donkey antibodies conjugated to Alexa 555 and goat antibodies conjugated to Alexa 488 (Molecular Probes) were used at 1:500.

### Fish husbandry and set up

Zebrafish were maintained within the *Danio rerio* University of Melbourne (DrUM) fish facility at the University of Melbourne (ethics approval ID10400, 22235), in accordance with local guidelines. All experimental procedures were approved by the Faculty of Science Animal Ethics Committee at the University of Melbourne (ethics approval ID10232). Adult zebrafish were kept at 28.5°C on 12/12h hour light/dark cycles. Zebrafish lines used were Tg(*lws2:nfsb-mCherry*), Tg(*gfap:eGFP*) (Raymond et al., 2006), Tg(*her4.1:dRFP*). For breeding, 2-4 males and females were set up in breeding tanks overnight, separated by a clear barrier, which was removed the next morning. Fertilized eggs were collected in E3 (5mM NaCl; 0.17mM KCl; 0.33mM CaCl_2_; 0.33mM MgSO_4_) at a maximum density of 50 eggs/40mL petri dish.

### Morpholino injection and electroporation

Protocol as per Thummel et al., 2011 (Thummel, Bailey et al. 2011). Morpholino (MO) oligos were custom-designed and ordered through GeneTools (Philomath, OR) and injected into adult (6-13 months) zebrafish retinas, followed by electroporation and recovery. Treatment fish were then immediately subjected to injury, controls were kept in fish water.

### Zebrafish injury model

Two transgenic lines used were crossed to create a metronidazole (MTZ) inducible photoreceptor ablation line. Tg(*lws2:nfsb-mCherry*), in which the long wavelength sensitive (*lws2*) promoter drove expression of the oxygen-insensitive NAD(P)H Nitroreductase (nfsB) gene, encoding the Nitroreductase (NTR) enzyme fused to the mCherry fluorescent reporter protein. NTR converted MTZ into a cytotoxin, causing DNA mutations and resulting in precise ablation of the targeted neurons (Curado, Stainier et al. 2008), which could be visualised by loss of the mCherry tag. In order to induce ablation adult fish were swum in 10mM MTZ in adult fish water for 24 hours at 28.5°C.

### Zebrafish drug treatments

Larvae were swum in 5% 1,6-Hexanediol (Sigma, #240117) or 2,5-Hexanediol (Sigma, #H11904) in fish water for two minutes at 6dpf, before returning to fish water.

### Husbandry of adult zebrafish in laboratory during experiments

Starting with the electroporation day, adult fish were not fed. After the procedure, fish were kept off constant water flow in 28.5°C incubators with 12h/12h light/dark cycle. They were monitored twice daily (including pH and ammonia test) and the water was changed daily for 1 – 5 days preceding their humane death.

### Immunostaining fish samples

Adult fish were humanely killed between 4-7 days post injury (dpi) by overdose with 1000 ppm AQUIS (Primo Aquaculture, #106036). Eyes were then dissected from the fish and fixed in 4% paraformaldehyde (PFA) in phosphate-buffered saline (PBS) at 4°C overnight. Samples were washed 3 times for 10 minutes in PBS, immersed in 30% sucrose for 24 hours at 4°C, and immersed in a 50/50 solution of Optimal Cutting Temperature (OCT) compound and 30% sucrose for 24 hours at 4°C. Eyes were embedded in 100% OCT, frozen into moulds using an ethanol/dry ice bath, and stored at −20°C.

Embedded tissue was sectioned using a Leica Cryostat (object temperature −20°C, chamber temperature −16°C). Transverse zebrafish retina sections were cut at 12μm thickness, collected onto Superfrost slides, and left at room temperature to dry before being stored at −20°C.

The sections were defrosted at room temperature before being washed for 10 minutes in PBS. Antigen retrieval was performed for PCNA staining by incubating slides in 2M HCl for 30m at 60°C. Antibodies were diluted in 5% fetal bovine serum (FBS) in PBS. The slides were covered with Parafilm and tissue was dampened with PBS, before placing at the bottom of the staining container to provide humidity. After overnight incubation at 4°C, the slides were washed 3 times for 10 minutes in PBS, the secondary antibodies were applied and left at room temperature for 3-4 hours. The sections were then washed 2 times with PBS for 10 minutes each to remove excess secondary antibody. A final 15-20 minute wash containing 100 μ/L 4,6-diamidino-2-phenylindole (DAPI) was performed to stain the nuclei, after which slides were cover-slipped with Mowiol, left to dry overnight at room temperature, and stored at 4°C. Primary antibodies used were mouse anti-Prox1 (1:2000-1:10000, Invitrogen), mouse anti-GS (1:500, Merck MAB302), rabbit anti-PCNA (1:400, Sigma SAB2701819), rabbit anti-Hes1 (1:500, Invitrogen #MA5-32258), mouse anti-Zpr1 (1:500, Zebrafish International Resource Centre), rabbit Anti-Hes4 (1: 500, Invitrogen #PA5-84551). Secondary goat antibody conjugated to anti-mouse or anti-rabbit Alexa 488 or 647 (Zebrafish International Resource Centre) were used at 1:500.

### Microscopy and analysis

Slides were imaged using the Olympus FV3000 confocal microscope within the Centre for Advanced Histology and Microscopy (CAHM) at the Peter MacCallum Cancer Centre. A 40x or 60x objective lens was used to capture images using 405nm, 488nm, 561nm, and 633nm channels. Images were processed with Fiji Is Just ImageJ (Fiji) and Photoshop.

### Statistical analysis

#### Analysis of clones in fly brains

At least eleven animals per genotype were used for all experiments. Volume of clones or regions of interest was estimated from three-dimensional reconstructions of 2 μm spaced confocal Z stacks with Volocity software (Improvision).

Clone volume was calculated by making a Region Of Interest (ROI) around the GFP+ clone (excluding the top section where the superficial NBs are). The rate of dedifferentiation was represented as the volume of Mira+ cells as a percentage of clone.

#### Analysis of zebrafish retinal sections

At least five animals per genotype were used for all experiments. All analysis was carried out on Fiji using freehand selections and measure tools.

Quantification of the percentage of PCNA+/Prox1+/Zpr1+ cells within the INL or ONL was calculated by manually drawing around the INL/ONL and measuring the area, followed by analysing the mean number of voxels in a given channel within that area. The percentage of PCNA+/Prox1+/Zpr1+/Hes1+ cells within the INL or ONL was therefore represented as the % of positive voxels within a given area. The number of PH3+ cells within the INL/ONL was calculated by manually counting the number of PH3+ cells and normalising it to the area of the INL/ONL.

In all graphs and histograms, error bars represent the standard error of the mean (SEM) and p values are calculated by performing two-tailed, unpaired Student’s t test. The Welch’s correction was applied in case of unequal variances.

#### Single-cell sample preparation

Zebrafish line Tg(*her4.1:dRFP/gfap:GFP/lws2:nfsb-mCherry*) for red cone ablation, was used in this study. Zebrafish larvae at 6 days post-fertilisation (dpf) were swum in a 10mM solution of metronidazole for 48 hours, to specifically eliminate the relevant subtype of cone photoreceptor. At 8dpf, fish were rinsed in fresh system water under standard housing conditions. At 9dpf (3dpi), fish were humanely killed. The single-cell suspensions of 9dpf zebrafish were prepared by following a published protocol (Lopez-Ramirez et al., 2016). Retinae were dissected and digested in 350 µl papain solution at 37°C for 15 minutes. The papain solution was prepared as follows: 100 µl papain (Worthington, LS003126), 100ul of 1% DNase (Sigma, DN25), and 200 µl of 12mg/ml L-cysteine (Sigma, C6852) were added to a 5 ml DMEM/F12 (Invitrogen, 11330032). During digestion, retinal tissue was mixed by pipetting up and down 4 x 10 times. Following digestion, 1400 µl of washing buffer was added, containing 65 µl of 45% glucose (Invitrogen, 04196545 SB), 50ul of 1M HEPES (Sigma, H4034), and 500 µl FBS (Gibco, 10270106) in 9.385ml of 1x DPBS (Invitrogen, 14190-144). All solutions were filtered through a 0.22 µm filter (Millipore) to sterilise and stored at 4°C prior to use.

#### 10X Chromium single-cell RNA sequencing

To perform single-cell RNA sequencing (scRNA-seq), FACS-isolated cells (sorted for *gfap:eGFP*) were loaded onto the Chromium Single Cell Chip (10x Genomics, USA) according to the manufacturer’s protocol. The scRNA-seq libraries were generated using the GemCode Single-Cell Instrument and Single Cell 3’ Library and Gel Bread kit v2 and v3 Chip kit (10x Genomics, 120237). Library quantification and quality assessments were performed by Qubit fluorometric assay (Invitrogen) and dsDNA High Sensitivity Assay Kit (AATI, DNF-474-0500). The fragment analyser was performed using the High Sensitivity Large Fragment −50kb Analysis Kit (AATI, DNF-464). The indexed library was tested for quality, and sequenced by the Illumina NovaSeq 6000 sequencer with the S2 flow cell using paired-end 150 x 150 base pairs as the sequencing mode. Sequencing depth was 60K reads per cell.

#### Quality filtering and pre-processing

Filtered matrix raw data files were further analysed in R using Seurat. Prior to downstream filtering, 10520 cells were obtained. Post-filtering and exclusion of non-Müller glia-derived cell types outputted achieved 9789. Low quality cells or cells containing doublets were excluded from all datasets; reads between 200 and 4500 genes per cell were included. Cells with a percentage of mitochondrial gene expression of greater than 35% were excluded from the analysis.

Samples were analysed using Seurat::NormalizeData, variable features for downstream analysis were identified using Seurat:: FindVariableFeatures, and scaled using Seurat::ScaleData. An optimal number of principal components (PCs) (generated through Seurat::RunPCA) for dimensional reduction were selected using the function Seurat::ElbowPlot. PCs containing the greatest variance were selected. Out of the 20 PCs originally specified, 15 PCs were selected to guide the unbiased clustering analysis. Clustering was performed using the shared nearest neighbour (SNN), graph based approach through Seurat::FindNeighbours. Seurat::FindClusters produced a uniform manifold approximation and projection (UMAP) plots containing 12 clusters.

## Results

### Candidate genes: *elav*, *dpn*, *pros*, and *SoxN* regulate dedifferentiation in the *Drosophila* medulla

Key transcription factors such as Nervous fingers-1 (Nerfin-1), Midlife Crisis (Mdlc), and Longitudinal lacking (Lola) were shown by us and others to maintain neuronal identity in *Drosophila* (Carney et al., 2013, Southall et al., 2014, Froldi et al., 2015, Vissers et al., 2018). In the medulla, the *Drosophila* visual processing centre, we previously showed that Nerfin-1 maintains neuronal fate in conjunction with its co-factor the Hippo pathway transcription factor, Scalloped (Sd) (Vissers et al., 2018). Interestingly, the vertebrate ortholog of Nerfin-1, Insulinoma-associated 1a (Insm1a), is a master regulator of zebrafish retinal regeneration, playing roles in multipotency and cell cycle exit (Ramachandran et al., 2012). Our targeted DamID analysis in the medulla neurons identified 3587 target genes commonly regulated by Nerfin-1 and Sd (Vissers et al., 2018). 470 candidate Sd/Nerfin-1 target genes were upregulated in *Drosophila* neural stem cells (neuroblasts, NBs) versus neurons, while 656 genes were downregulated. 36 of these genes encode transcription factors, of which, 12 were expressed in either NBs or neurons. To assess if any of these genes were functionally involved in dedifferentiation, we used a heat shock-induced *actin-GAL4 flp-out* (*hs flp*) (Figure 1D), to induce clones that either overexpress or knockdown candidate genes. RNAis were used to knockdown the expression of these genes if they were expressed in neurons, and UASs were used to overexpress the candidate genes expressed in NBs. Additionally, we used a medulla driver (*GMRH108-GAL4*) to validate our findings (the result of our dedifferentiation screen is summarised in Figure S1).

**Figure 1.**
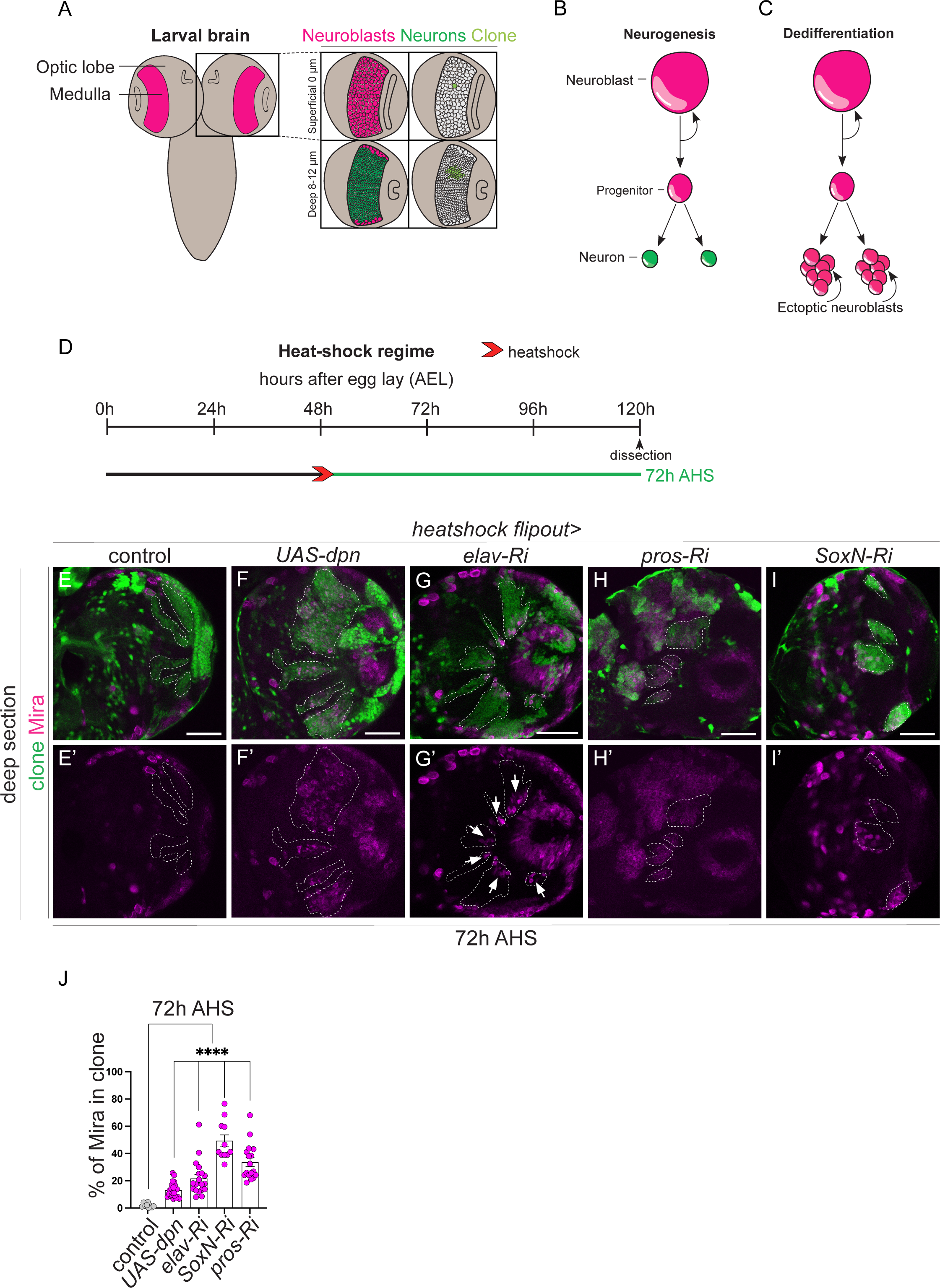
Knockdown of Dpn, Elav, Pros, and SoxN result in dedifferentiation of medulla neurons. (A) Schematic representation of the larval central nervous system (CNS). Medulla (depicted in pink) is located with the Optic Lobes (OLs) of the CNS. Within boxes: (left) Superficial surface (0μm) is occupied by neuroblasts (NBs, pink) and deep layers (8-120μm) are occupied by neurons (green), (right) Cross section through a hs flp clone (light green) showing the clone consists of a single NB in the superficial layer and its neuronal progeny in the deep layer. (B) Schematic representation of NB divisions: NBs undergo asymmetric division to give rise to a Ganglion mother cell (GMC) which produces two neurons. (C) Schematic representation of dedifferentiation: after neurons dedifferentiate, ectopic neuroblasts divide and create tumours. (D) Schematic depicting the heat shock regimes used in (E-I’). Clones were induced via heat shock (red arrows), and dissected at 72 hr (green) after heat shock. (E-I’) Representative images of the deep medulla neuronal layer in the larval OL, in which *UAS-dpn*, *elav-Ri*, *pros-Ri*, *SoxN-Ri*, and control are driven in clones by *hs flp* (marked by GFP and outlined) and stained with the stem cell marker, Miranda (Mira, magenta). At 72h after clone induction, numerous Mira+ NBs are recovered in *UAS-dpn*, *elav-Ri*, *pros-Ri*, and *SoxN-Ri* clones but not in control clones. Quantified in J. (J) Quantification of % Mira+ cells in *UAS-dpn*, *elav-Ri*, *pros-Ri*, *SoxN-Ri*, and control clones (calculated as the ratio of Mira+ cell volume as a percentage of total clone volume). Control n=9, m=1.874 ±0.4897, *UAS-dpn* n=17, m=33.62 ±3.243, *elav-Ri* n=22, m=13.16 ±1.164, *SoxN-Ri* n=19, m=21.74 ±2.926, *pros-Ri* n=11, m=49.34 ±4.36. Data are represented as mean ± SEM. P-values were obtained using unpaired t-test, and Welch’s correction was applied in case of unequal variances. ****p < 0.0001. Scale bars: 50 μm. Deadpan data is adapted from Veen et al., under revision (Veen, 2022).

As medulla NBs are only observed in the superficial layer and neurons in the deep layers, we scored for dedifferentiation by assessing the presence of the NB marker, Miranda (Mira), in the deep neuronal layers of the medulla (Figure 1A). We hypothesised that candidate transcription factors specifically expressed in the neurons would act as transcriptional repressors to silence the neural stem cell fate. Therefore, downregulating these genes in the neurons, should yield ectopic NBs, indicative of dedifferentiation (Figure 1B). Conversely, transcription factors expressed in the NBs would act as transcriptional activators, and must be silenced in the neurons to maintain differentiation. Overexpressing these factors in the neurons would produce ectopic NBs. Upon induction of the bHLH transcription factor, Deadpan (Dpn), we observed ectopic NBs (as indicated by Mira+ cells) in the medulla neuronal layer of the larval CNS compared to control clones (Figure 1E-F’, J). Similarly, knockdown of the homeobox transcription factor, Prospero (Pros) and high mobility group box (HMGB) transcription factor, SoxNeuro (SoxN), induced numerous ectopic Mira+ cells in the deep layer (Figure 1 H-I’, J). Interestingly, knockdown of Embryonic lethal abnormal vision (Elav) resulted in the formation of ectopic NBs only in a subset of medulla neurons (Figure 1 G-G’, arrows, J). Therefore, Elav may only be required to maintain the identity of a subset of neurons in a temporal fashion. Together, this data shows that overexpression of Dpn and knockdown of Pros, Elav, and SoxN phenocopied Nerfin-1 loss of function, and thus are likely mediators of dedifferentiation. As factors implicated in dedifferentiation may be involved in the regenerative response upon injury, we next examined whether these factors are involved in zebrafish retina regeneration.

### Characterisation of MTZ-induced adult photoreceptor injury model

To study the process of regeneration in adult zebrafish, we used a metronidazole (MTZ) inducible transgenic fish line to specifically ablate the highly abundant long wavelength sensitive (lws) PRs. As previously described (Montgomery et al., 2010), adult fish between the ages of 5 and 11 months of age, were subject to MTZ for 24 hours, followed by a 72h recovery time. We observed similar amounts of PR reduction (mCherry-labelled cells) at 72 hours post injury (hpi) in fish that were either 5 or 11 months old (Figure S2A-D’’’). This suggests that MTZ-induced injury was similarly effective for fish of either age, therefore, for the rest of the experiments, fish between 10 and 12 months were utilized.

Next, we conducted a time course experiment at 16, 24, 48, 72, and 96 hpi to characterise the regenerative response under our ablation paradigm. Müller glia (MG), labelled by Glutamine Synthase (GS), are known to produce progenitor cells, labelled by Proliferating Cell Nuclear Antigen (PCNA) upon injury (Figure S3A). Consistent with other adult injury paradigms (Kassen et al., 2007), MG-derived progenitors were first observed at around 24hpi (Figure S3 D-D’’’). They began proliferating at 48hpi, as indicated by clusters of PCNA+ cells across the inner nuclear cell layer (INL), observed by 72hpi (Figure S3 E-E’’’, outlined). This increase in PCNA+ cells was accompanied by a more diffuse GS staining pattern at 72 and 96hpi, consistent with the dilution of MG markers into the neuronal progenitors (Figure S3 F’’-F’’’, G’’’, outlined). The regenerative response observed in our MTZ induced injury model is comparable to that reported for other PR injury paradigms (Kassen et al., 2007), therefore, our MTZ model of degeneration is consistent with other previously described damage models.

### Her6/Hes1 is downregulated in MG-derived progenitors

As the *Drosophila* genes *deadpan* (*dpn*) and *prospero* (*pros*) induced dedifferentiation (Figure S1) and their homologs *hairy-related 6* (*her6*) and *prospero homeobox 1* (*prox1*) have not previously been implicated in PR regeneration, we next set out to assess the function of these novel regulators utilising our MTZ-mediated injury model.

Dpn is expressed in NBs and necessary for the maintenance of NB ‘stemness’ (San-Juán annbd Baonza 2011). The zebrafish ortholog of *dpn*, *her6*, is expressed in developmental progenitors within the inner nuclear layer (INL) and is downregulated upon neurogenesis (Soto et al., 2020). To investigate whether Her6 played a role during regeneration, we assessed its expression in a time course analysis upon PR injury, between 16 and 96hpi (Figure 2B) using an antibody against the murine ortholog of Her6, Hairy and enhancer of split-1 (Hes1). We found that Hes1 was expressed in the INL (Figure 2, C-C’’, outlined) in the control retina. Upon injury, between 16 and 24hpi, Hes1 expression appeared downregulated, and by 48hpi Hes1 was no longer detected within the PCNA+ progenitor cells (Figure 2 C’-H’’). This suggests that Hes1 is downregulated upon progenitor formation, and its downregulation may be required for the proliferative response in the context of regeneration.

**Figure 2.**
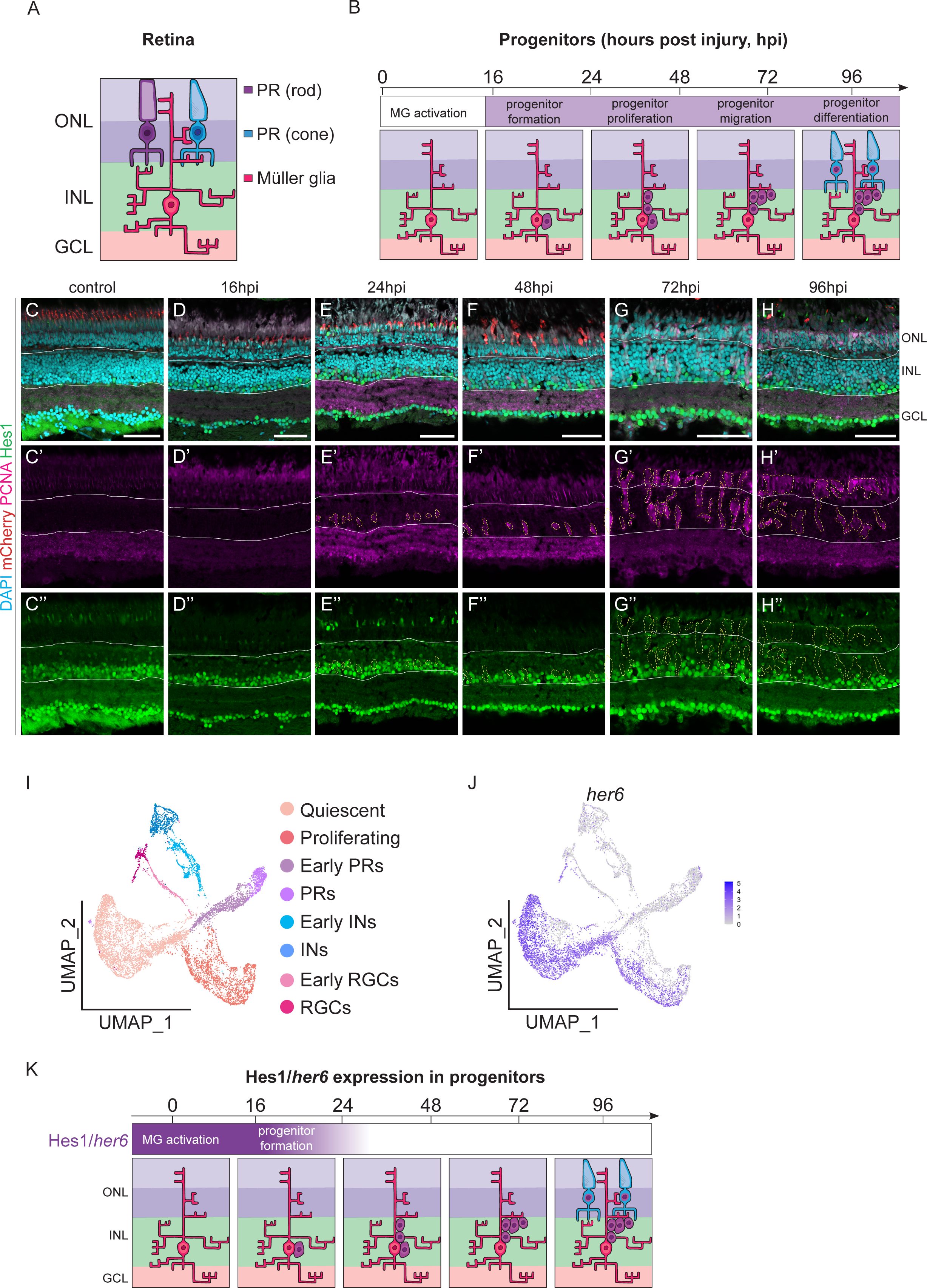
Her6/Hes1 is downregulated in MG-derived progenitors. (A) Schematic representation of retinal cells. Rod and cone photoreceptors (blue and purple, respectively) are located in the outer nuclear layer (ONL). Müller glia are located within the inner nuclear layer (INL). Ganglion cell layer (GCL) is represented in pink. (B) Schematic representation of the Müller glia-derived progenitors in the regenerating retina. Time course from 0-96 hours post injury (hpi). A Müller glia (pink, MG) derived progenitor is generated (purple, second panel); the progenitors proliferate (third panel) and migrate (fourth panel) then differentiate into PRs (fifth panel). (C-H’’) Micrographs of retinal sections with the Inner Nuclear Layer (INL) outlined. Ablation of PRs (red) was induced with metronidazole (MTZ) treatment in Tg(*lws2:nfsb-mCherry*) zebrafish. Sections were stained with Proliferating Cell Nuclear Antigen (PCNA, pink) to label proliferating cells, Hairy-enhancer of split 1 (Hes1, green), and DAPI to label nuclei. Within the control, 16hpi, and 24hpi Hes1 was expressed in the lower half of the INL (outlined) (B-D’’). From 48hpi to 96hpi Hes1 was not expressed in PCNA+ cells within the INL (E-G’’). Scale bars: 50 μm. (I) UMAP plot of FACS MG 3 days after injury outlining different MG populations: Quiescent (beige), Proliferating (coral), Early PRs (light purple), PRs (purple), Early Inhibitory neurons (INs, light blue), INs (blue), Early Retinal ganglion cells (RGCs, light pink), RGCs (pink). (J) Feature plot that shows her6 was expressed mainly in quiescent and some proliferating MG populations and downregulated in differentiating cell populations. (K) Schematic representation of Hes1/*her6* expression during regeneration. Hes1/*her6* is present in the initial progenitors and downregulated upon their proliferation.

To further assess *her6* expression, we turned to a larval model. Firstly, we performed an injury time course looking at PCNA and GS, identical to what we performed in adults, to confirm its similarity to the adult injury paradigm. We found that, MG-derived progenitors first appeared by 24-48hpi and began proliferating by 72hpi (Figure S4), validating the resemblance between adult and larval injury models. As larval and adult regenerative response during injury is comparable, we assessed the expression of *her6* in a single-cell RNA-seq dataset from a larval MTZ-induced injury model. MG from larval zebrafish were sorted and sequenced at 72hpi. Through unbiased UMAP clustering, we characterised a heterogenous population of differing MG cell states using key known markers in our previous study (Krylov et al., currently under revision). These consist of quiescent and activated MG as well as proliferating neuronal progenitors and differentiating MG-derived progenitor cells: early PRs, PRs, early inhibitory neurons (INs), INs, early retinal ganglion cells (RGCs), and RGCs (Figure 2A,I). We found that *her6* was highly expressed within cells in the quiescent and reprogrammed MG and was downregulated in the proliferating and differentiating populations of neuronal progenitors (Figure 2 I-J). This is consistent with our time-course analysis in the adult injury model, where we observed a downregulation of Her6 in the PCNA+ MG progenitor cells (Figure 2K). Together, our data suggest that Her6 may act as in inhibitory regulator, and its downregulation may be required for regeneration to occur.

### The loss of Her6 results in an increase of PRs after injury

To further explore Her6 during regeneration, we assessed whether it was functionally required. We knocked down *her6* expression via *in vivo* electroporation of morpholino (MO) within the retina (Thummel et al., 2011) and successfully reduced Her6 expression (Figure S5 A-D’). We observed no significant change in the number of PCNA+ progenitor cells at 48 and 72hpi, compared to the standard control (SC) MO (Figure 3 A-E). However, we observed an overall increase of the number of cones, indicated by the ZPR1 Zinc Finger (Zpr1) antibody (which labels Arrestin 3 (Arr3) expression (Ile et al., 2010)) at 72hpi (Figure 3 F-G’’, H), indicating that Her6 knockdown has increased the overall production of PRs. As the number of progenitors did not change, it is likely that cell cycle speed of progenitor cells may be elevated. To test this, we assessed the amount of Phospho-Histone3 (PH3) within the INL and ONL, and found that there was a higher number of PH3+ cells in the *her6* MO-treated fish at 72hpi, compared to control (Figure 3I-M). This suggests that the loss of Her6 increases the speed at which progenitors progress through the cell cycle. Together, our data suggests that Her6 is a negative regulator of regeneration and its downregulation promotes PR production upon injury (Figure 3N).

**Figure 3.**
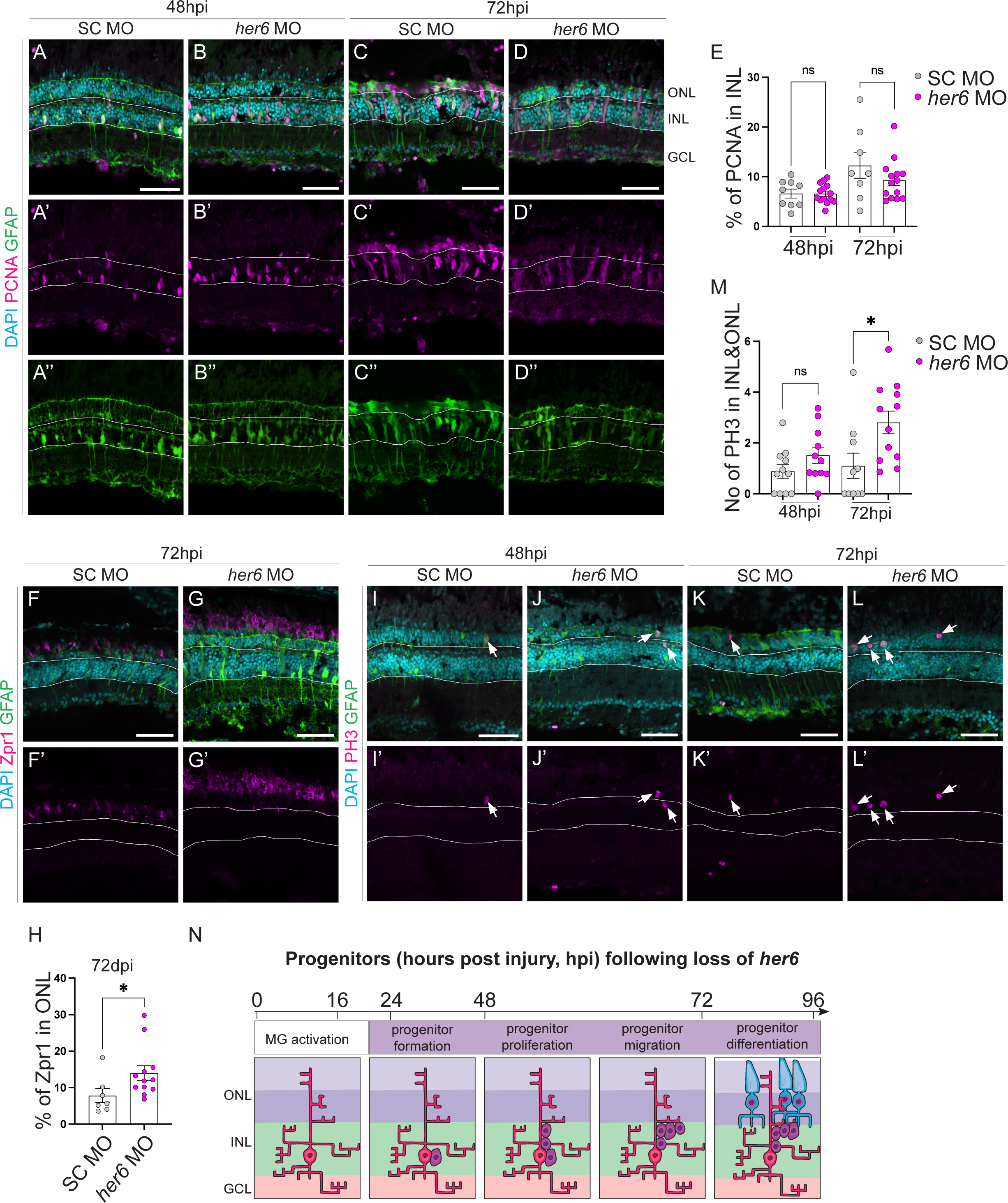
Loss of Her6 increases PR production after ablation. (A-D’’, F-G’, I-L’) Micrographs of retinal sections comparing regeneration in control and Her6 loss of function. Tg(*lws2:nfsb-mCherry, gfap:eGFP*) zebrafish were injected with either a standard control (SC) or *hairy-related 6* (*her6*) morpholino (MO) prior to ablation. Sections were stained with Proliferating Cell Nuclear Antigen (PCNA, pink) to label proliferating cells, ZPR1 Zinc Finger (Zpr1, pink) to label PRs, or Phospho-Histone3 (PH3, pink) to label mitotic cells, and DAPI to label nuclei. GFAP (green) labels MG. Compared to SC MO, *her6* MO treated fish had similar expressions of PCNA and GFAP at 48hpi and 72hpi (A-D’’), quantified in E, higher expression of Zpr1 at 72hpi (F-G’’), quantified in H, similar expression of PH3 at 48hpi, and higher expression of PH3 by 72hpi, quantified in M. Scale bars: 50 μm. (E) Quantification of percentage of PCNA+ cells within the INL (calculated as the percentage of PCNA+ cells within the INL normalised by the area of the INL). 48hpi (SC MO n=9, m=6.621 ±0.9034, *her6* MO n=14, m=6.59 ±0.5219), 72hpi (SC MO n=8, m=12.27 ±2.579, *her6* MO n=14, m=9.299 ±1.1). (H) Quantification of percentage of Zpr1+ cells within the ONL (calculated as the percentage of PCNA+ cells within the INL normalised by the area of the ONL). SC MO n=7, m=7.809 ±1.938, *her6* MO n=12, m=13.97 ±2.202 (M) Quantification of percentage of PH3+ cells within the INL/ONL (calculated as the number of PH3+ cells within the INL/ONL normalised by the area of the INL/ONL). 48hpi (SC MO n=11, m=0.8812 ±0.2671, *her6* MO n=11, m=1.519 ±0.3166), 72hpi (SC MO n=10, m=1.102 ±0.4956, *her6* MO n=12, m=2.806 ±0.4442). (N) Schematic representation of progenitors following the loss of *her6*. Progenitor formation, proliferation, and differentiation occurs rapidly. Data are represented as mean ± SEM. P-values were obtained using unpaired t-test, and Welch’s correction was applied in case of unequal variances. *p < 0.05.

### Prox1 is expressed in differentiating PRs after injury

Next, we assessed whether the homolog of *Drosophila* Prospero (Pros), Prospero homeobox1a (Prox1), was involved in zebrafish retina regeneration. *Drosophila* Pros is required for cell cycle exit in NBs, and its loss results in the formation of ectopic stem cells (Betschinger et al., 2006). Prox1 is involved in the differentiation of inhibitory neurons (amacrine and horizontal cells) within the developing mouse retina (Dyer et al., 2003), however, the role of Prox1 in retinal regeneration is so far unknown.

To understand at what stage of regeneration Prox1 may be involved, we performed a time course analysis to assess Prox1 expression during regeneration. We found that Prox1 was consistently expressed within the INL of control retinas (Figure 4 A-F’’). Its expression was not altered in the MG-derived progenitors upon injury (Figure 4 C’’-F’’, outlined). However, Prox1 was upregulated in the differentiating PRs beginning from 72hpi (Figure 4 E’’-F’’).

**Figure 4.**
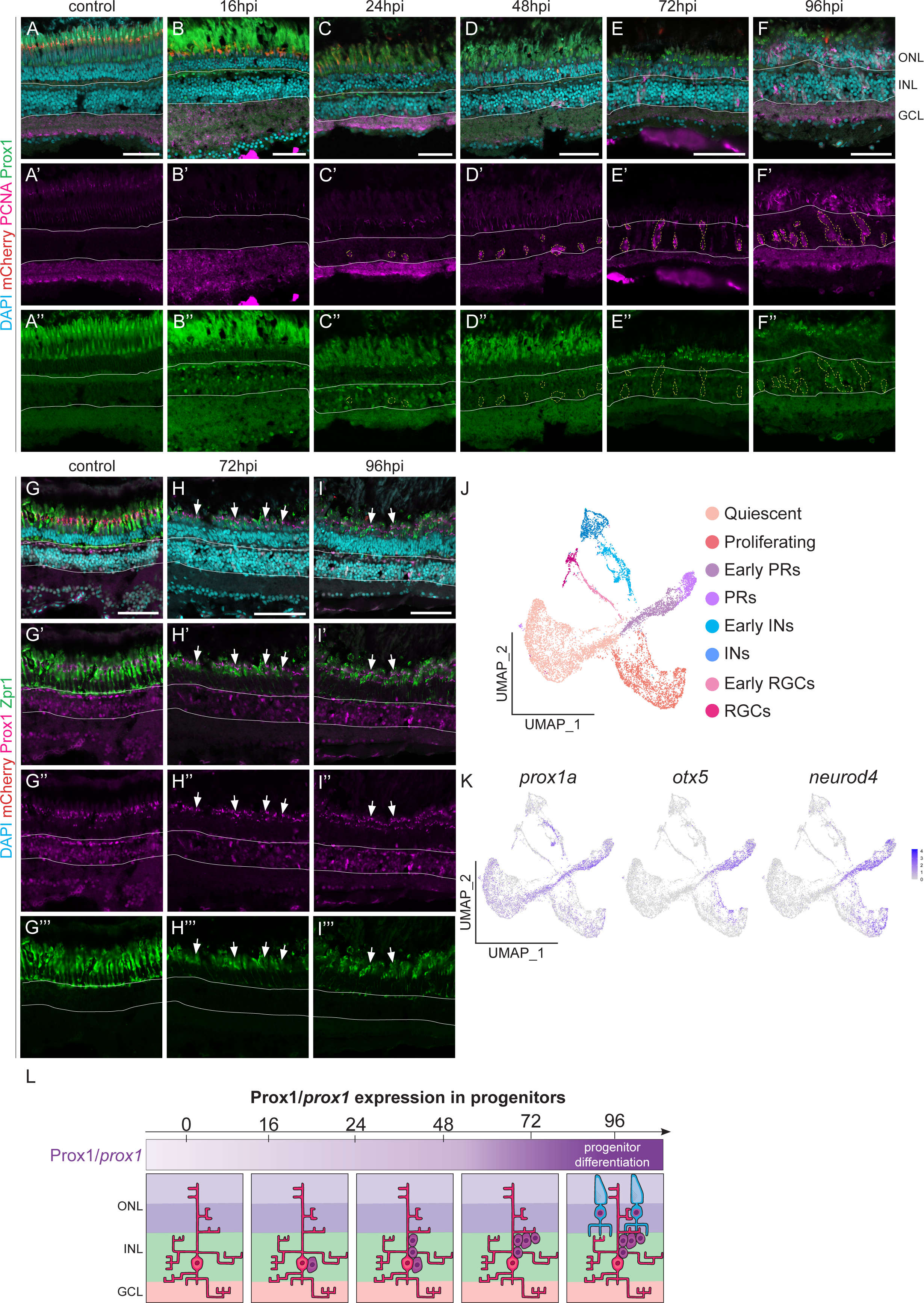
Prox1 is expressed in photoreceptors within the ONL. (A-I’’) Micrographs of retinal sections with the Inner Nuclear Layer (INL) outlined. Ablation of PRs (red) was induced with metronidazole (MTZ) treatment in Tg(*lws2:nfsb-mCherry*) zebrafish. Sections were stained with Proliferating Cell Nuclear Antigen (PCNA, pink) to label proliferating cells, and Prospero homeobox 1 (Prox1, green), ZPR1 Zinc Finger (Zpr1, green), and DAPI to label nuclei. (A-F’’) Within the control, 16hpi, 24hpi, and 48hpi Prox1 was expressed in the INL (solid outline) and not in PCNA+ cells (dotted outline) (B-E’’). At 72hpi and 96hpi Prox1 was expressed in cells within the INL and in the ONL, but not expressed in PCNA+ cells within the INL. (G-I’’) Zpr1 was expressed in the ONL in the control and all time points. At 72hpi and 96hpi Prox1 was expressed within Zpr1 expressing cells (arrows). Scale bars: 50 μm. (J) UMAP plot of FACS MG 3 days after injury outlining different MG populations: Quiescent (beige), Proliferating (coral), Early PRs (light purple), PRs (purple), Early Inhibitory neurons (INs, light blue), INs (blue), Early Retinal ganglion cells (RGCs, light pink), RGCs (pink). (K) Feature plot that shows *prox1a* was expressed in early PRs, characterised by the PR-specific genes *otx5* and *neurod4*. (K) Schematic representation of Prox1/*prox1* expression during regeneration. Prox1/*prox1* is present in differentiating PRs.

Consistent with this, we identified prominent *prox1a* expression within the differentiating PR population in our sc-RNAseq dataset, as *prox1a* is highly upregulated in the cell population where PR-specific genes, *neurod4* and *otx5* were also upregulated (Figure 4 J-K). To assess this qualitatively, we conducted an additional time course and studied Prox1 expression along with the PR-specific marker, Zpr1, at later timepoints (72 and 96hpi). Indeed, we found that Prox1 was expressed within Zpr1 expressing cells (Figure 4 G-I’’’, arrows). As Prox1 is expressed in differentiating PRs (Figure 4L), it is likely that it is required to promote PR differentiation after injury.

### Prox1 loss results in a reduction of PR production after injury

To assess the function of Prox1 in PR regeneration, we knocked it down via *in vivo* MO electroporation. We found at 72 and 120hpi, there was no significant change in the percentage of PCNA+ cells (Figure 5 A-E), suggesting Prox1 does not regulate progenitor formation or proliferation. However, there was a significant decrease in the percentage of cells expressing the mature PR marker, Zpr1, by 120hpi (Figure 5 F-J). Our data suggests that Prox1 is required for PR differentiation following injury, in its absence, differentiated mature PRs fail to be produced (Figure 5 K).

**Figure 5.**
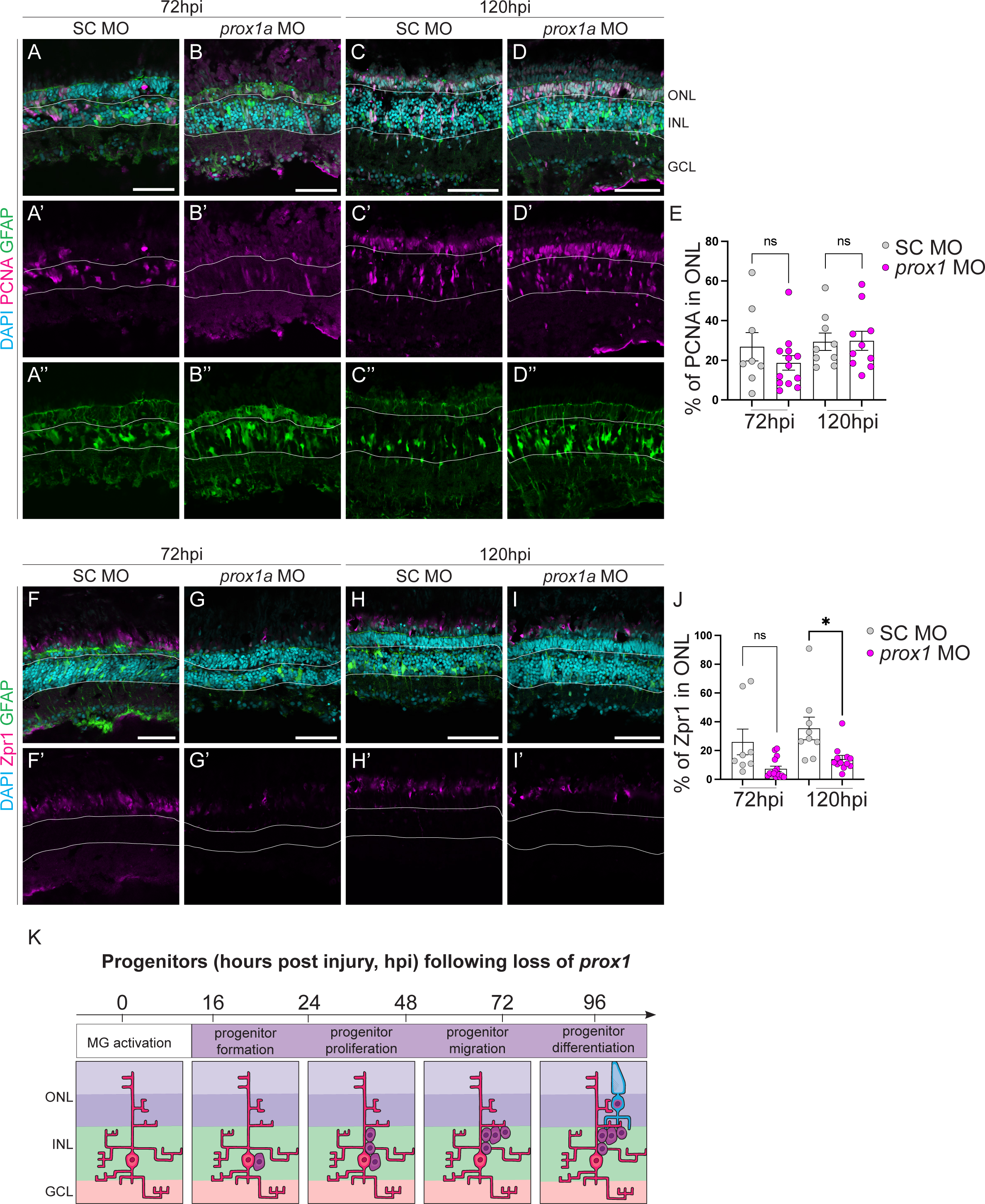
Loss of Prox1 decreases PR production after injury. (A-D’’, F-I’) Micrographs of retinal sections comparing regeneration in control and Her6 loss of function. Tg(*lws2:nfsb-mCherry, gfap:eGFP*) zebrafish were injected with either a standard control (SC) or *prospero homeobox 1a* (*prox1a*) morpholino (MO) prior to ablation. Sections were stained with Proliferating Cell Nuclear Antigen (PCNA, pink) to label proliferating cells, or ZPR1 Zinc Finger (Zpr1, pink) to label PRs, and DAPI to label nuclei. GFAP (green) labels MG. Compared to SC MO, *prox1a* MO treated fish had similar expressions of PCNA and GFAP at 72hpi and 120hpi (B-E’’, I). Quantified in (J). Compared to SC MO, *prox1a* MO treated fish had a lower expression of Zpr1 at 72hpi and 120hpi (F-G’’). Quantified in (K). Scale bars: 50 μm. (E) Quantification of percentage of PCNA+ cells within the INL (calculated as the percentage of PCNA+ cells within the INL normalised by the area of the INL). 72hpi (SC MO n=8, m=26.82 ±7.14, *prox1a* MO n=13, m=18.7 ±03.692), 120hpi (SC MO n=9, m=29.37 ±4.408, *prox1a* MO n=10, m=29.84 ±4.805). (J) Quantification of percentage of Zpr1+ cells within the ONL (calculated as the percentage of Zpr1+ cells within the ONL normalised by the area of the ONL). 72hpi (SC MO n=8, m=25.98 ±8.962, *prox1a* MO n=15, m=7.312 ±1.806, 120hpi (SC MO n=9, m=35.4 ±7.836, *prox1a* MO n=12, m=14.05 ±2.539). (K) Schematic representation of progenitors following the loss of *prox1a*; less PRs are generated. Data are represented as mean ± SEM. P-values were obtained using unpaired t-test, and Welch’s correction was applied in case of unequal variances. *p < 0.05.

### Prox1 may act through LLPS in PR differentiation

We next assessed the role of Prox1 in PR differentiation after injury. Interestingly, throughout our investigations Prox1 protein appeared in small puncta within the PRs (Figure 4 A-I’’’). This was reminiscent of the protein puncta the *Drosophila* ortholog of Prox1, Pros, forms during differentiation. Pros was shown to regulate neuronal differentiation though Liquid-liquid Phase Separation (LLPS) (Liu et al., 2020). We therefore assessed whether Prox1 drove PR differentiation during regeneration in a similar fashion. Many publications, including those using zebrafish models, assessed LLPS through 1,6-Hexanediol (1,6-HD) treatment (Krishnakumar et al., 2018). 1,6-HD is an organic compound commonly used to dissolve liquid-liquid phase separates (Duster et al., 2021). However, this has only been carried out *in vitro*. As it is more challenging to perform high throughput analysis in adult fish, we switched to the larval injury model (Figure S4) to first determine at which concentration 1,6-HD and the control 2,5-HD can be administered to zebrafish. We found that the treatment was the most effective and consistent when administered at 5% for 2 minutes (Figure 6B). Therefore, at 3dpf, we swam larvae in MTZ for 24 hours, allowed them to recover for 72 hours, then treated with either 1,6-HD or 2,5-HD for 2 minutes, and fixed and stained the larvae 24 hours later (at 96hpi). Overall, we found significantly less Prox1 in the ONL in 1,6-HD treated larvae compared to 2,5-HD treated larvae (Figure 6 C-D’’, G), suggesting that inhibiting phase separation may disrupt Prox1 levels. To assess whether phase separated Prox1 is functionally required for the regenerative response after injury, we examined Zpr1 expression in 1,6-HD treated larvae. We discovered that there was significantly less Zpr1 within the ONL in 1,6-HD treated larvae compared to control (Figure 6 D E’’’’, H). Together, our data demonstrates that Prox1 forms liquid-liquid phase condensates within PRs and this process is necessary for PR differentiation after injury.

**Figure 6.**
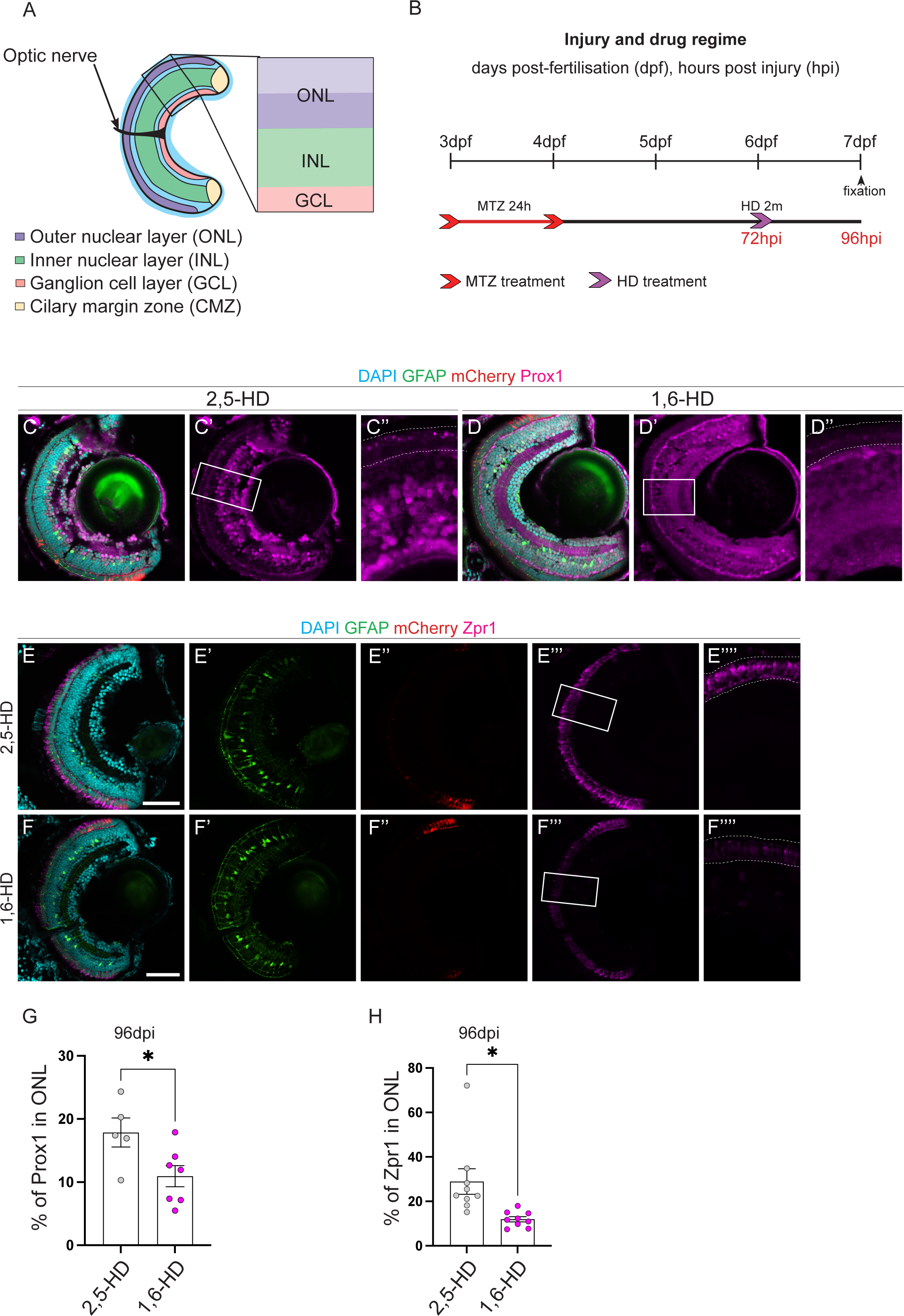
Prox1 regulates PR differentiation through LLPS. (A) Schematic representation of the larval retina labelling outer nuclear layer (ONL, purple), inner nuclear layer (INL, green), ganglion cell layer (GCL, pink), ciliary margin zone (CMZ, yellow) and optic nerve (arrow). (B) Schematic representation of injury and drug regime. Larvae were swum in metronidazole (MTZ) for 24 hours at 3dpf, followed by a 2m treatment with 1,6-Hexanediol (1,6-HD) or 2,5-HD at 6dpf, before their death and fixation. (C-F’’’) Micrographs of retinal sections. Following PR ablation at 3 days post fertilisation (dpf), Tg(*lws2:nfsb-mCherry, gfap:eGFP*) zebrafish were treated with 5% 1,6-HD or 2,5-HD for 2m. Sections were stained with Prospero homeobox 1 (Prox1, pink) or ZPR1 Zinc Finger (Zpr1, pink) to label PRs. GFAP (green) labels MG and DAPI labels nuclei. 1,6-HD treated larvae had reduced Prox1 expression in the ONL compared to 2,5-HD treated larvae. (C’’) and (D’’) represent boxes in (C’) and (D’), respectively, ONL is outlined (C’’and D’’. Quantified in (G). 1,6-HD treated larvae had reduced Zpr1 expression in the ONL compared to 2,5-HD treated larvae (E-F’’’’). (E’’’’) and (F’’’’) represent boxes in (E’’’) and (F’’’), respectively, ONL is outlined. GFAP and mCherry expression is similar between 1,6-HD and 2,5-HD treated larvae. Quantified in (H). Scale bars: 50 μm. (G) Quantification of percentage of Prox1+ cells within the INL (calculated as the percentage of Prox1+ cells within the ONL normalised by the area of the ONL). 1,6-HD n=7, m=10.94 ±1.681, 2,5-HD n=5, m=17.86 ±2.3. (H) Quantification of percentage of Zpr1+ cells within the INL (calculated as the percentage of Zpr1+ cells within the ONL normalised by the area of the ONL). 1,6-HD n=9, m=11.95 ±1.16, 2,5-HD n=9, m=28.91 ±5.72. Data are represented as mean ± SEM. P-values were obtained using unpaired t-test, and Welch’s correction was applied in case of unequal variances. *p < 0.05.

### Loss of *her6* increases Prox1 expression

As Her6 and Prox1 knockdowns both affected PR production during regeneration, it is interesting to speculate whether they function in the same pathway. DNA adenine methyltransferase IDentification (DamID) analysis, a tool mapping binding sites of chromatin binding proteins, previously showed the *Drosophila* ortholog of *prox1*, *pros*, is a target gene of the Her6 ortholog, Dpn (Southall and Brand, 2009). Additionally, Dpn acts in neural stem cells to repress Pros, and its loss results in increased Pros expression and differentiation (Zhu et al., 2012). In the regeneration context, we showed that Her6 knockdown is required for enhanced progenitor proliferation, resulting in the generation of additional PRs, and Prox1 is a regulator of PR differentiation. To examine whether Her6 and Prox1 intersect, we assessed Prox1 expression following Her6 knockdown via MO. Interestingly, at 48hpi there was significantly more Prox1 expression compared to control (Figure 7 A-E). 48hpi is an early time point during regeneration, further suggesting that the loss of Her6 may relieve the suppression of Prox1 at an earlier time point, to promote precocious PR generation. However, by 72hpi, the number of Prox1 cells is no longer significantly different between *her6* MO and control. Together, this data suggests that Prox1 may lie downstream of Her6 and is negatively regulated by Her6 (Figure 7K).

**Figure 7.**
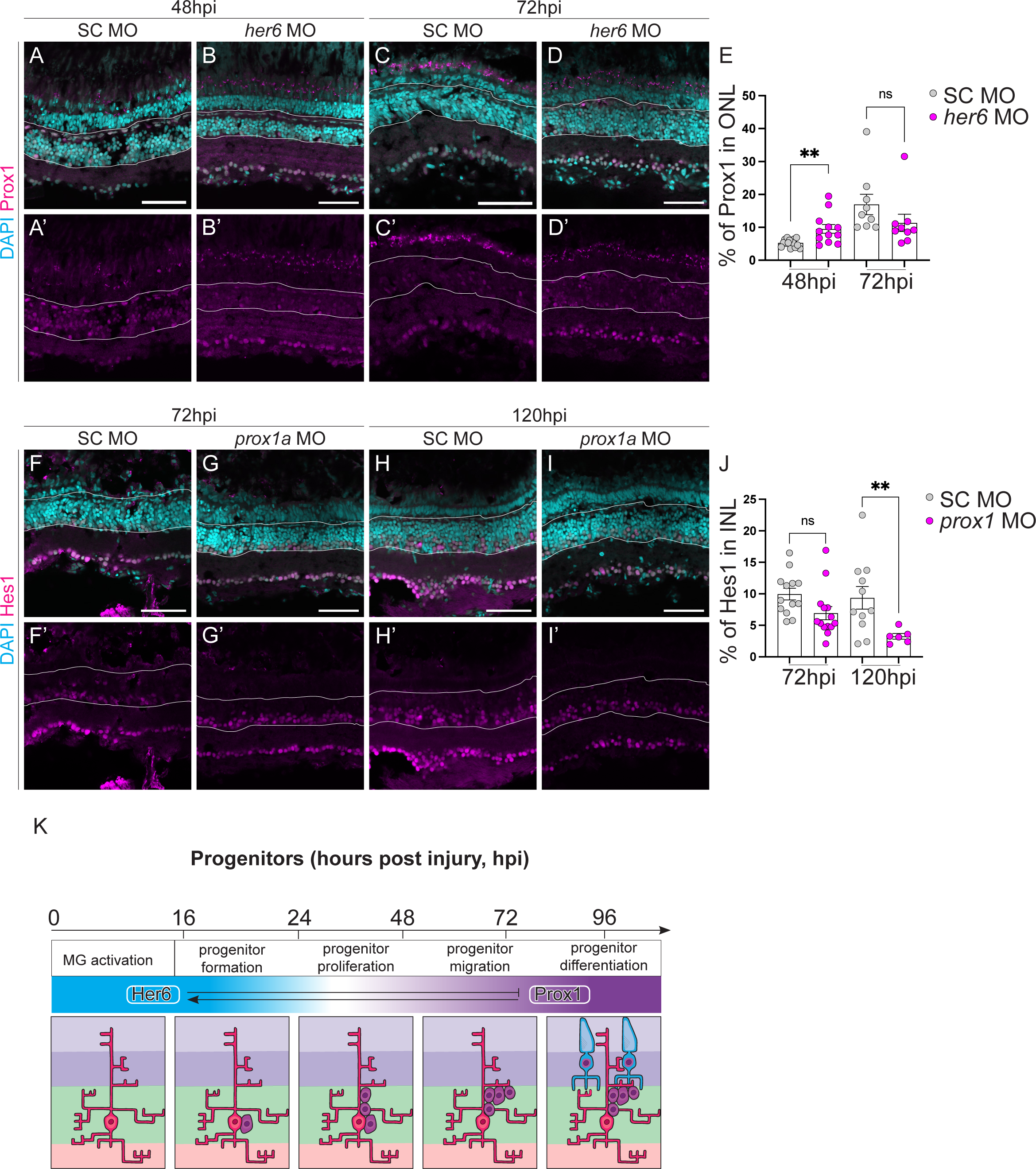
Her6 reduction increases Prox1 expression and subsequent PR regeneration. (A-D’’, F-I’) Micrographs of retinal sections comparing regeneration in control and Her6 loss of function. Tg(*lws2:nfsb-mCherry, gfap:eGFP*) zebrafish were injected with either a standard control (SC), *prospero homeobox 1a* (*prox1a*) morpholino (MO), or *hairy-related 6* (*her6*) MO prior to ablation. Sections were stained with Prox1 (pink) or Hes1 (pink), and DAPI to label nuclei. Compared to SC MO, *her6* MO treated fish had less Prox1 expression at 48hpi and a similar expression at 72hpi (A-D’), quantified in E. Compared to SC MO, *prox1a* MO treated fish had less Hes1 expression at 120hpi and a similar expression at 72hpi (A-D’), quantified in J. Scale bars: 50 μm. (E) Quantification of percentage of Prox1+ cells within the ONL (calculated as the percentage of Prox1+ cells within the ONL normalised by the area of the ONL). 48hpi (SC MO n=15, m=5.294 ±0.3032, *her6* MO n=12, m=9.507±1.339, 120hpi (SC MO n=9, m=16.97 ±3.096, *her6* MO n=9, m=11.34 ±2.625). (J) Quantification of percentage of Hes1+ cells within the INL (calculated as the percentage of Hes1+ cells within the INL normalised by the area of the INL). 72hpi (SC MO n=13, m=9.958 ±0.9212, *prox1a* MO n=14, m=6.929±1.044, 120hpi (SC MO n=11, m=9.355 ±1.792, *prox1a* MO n=6, m=3.255 ±0.4484). (K) Schematic representation of Her6 and Prox1expression during regeneration. Her6 is expressed during progenitor formation while Prox1 is present in differentiating PRs. Her6 is a negative regulator of Prox1 and Prox1 is a positive regulator of Her6. Data are represented as mean ± SEM. P-values were obtained using unpaired t-test, and Welch’s correction was applied in case of unequal variances. *p < 0.05.

Finally, we examined whether Her6 is also regulated by Prox1. Upon knockdown of Prox1, we found that Hes1 levels were significantly reduced (Figure 7 F-J). Therefore, in addition to the role of Prox1 in regulating PR differentiation, Prox1 is required to positively promote Her6 levels (Figure 7K).

Together, through examining the role of factors implicated in dedifferentiation in *Drosophila* optic lobes, we have identified two novel regulators of zebrafish retinal regeneration. Her6 negatively inhibits PR production via regulating the rate of the cell cycle progression of the progenitor cells, and the onset of Prox1. Prox1 mainly regulates PR differentiation, to promote the overall production of Zpr1+ PR cells.

## Discussion

In this study, we identified two evolutionary highly conserved novel candidates of regeneration, allowing us to identify pro-regenerative mechanisms, which can be used to improve the human regenerative response following injury. Indeed, many of the genes already identified to positively regulate PR regeneration in zebrafish were shown to improve neural regeneration in the mammalian retina (Todd et al., 2021, Jorstad et al., 2017, Pollak et al., 2013). This highlights the necessity and utility of such cross-species research.

### Her6 and Notch signalling in regeneration

We found that Her6 is downregulated upon progenitor formation and proliferation; furthermore, downregulation of Her6 enhances PR production. This suggests that Her6 is a negative regulator of regeneration, and it acts specifically in early progenitor cells. Interestingly, *her6* and its human ortholog, *hairy and enhancer of split 1* (*hes1*), are well-defined Notch targets (Pasini et al., 2004, Moriyama et al., 2006, Zhang et al., 2010). Similar to Her6, Notch signalling is downregulated during retinal regeneration to enable Müller glia (MG) to enter the cell cycle (Campbell et al., 2022, Fogerty et al., 2022). Additionally, the loss of Notch signalling is sufficient to induce MG proliferation in the undamaged retina (Conner et al., 2014). It would therefore be interesting to test whether the loss of Her6 can stimulate quiescent MG to re-enter cell cycle in the absence injury. Although the *Drosophila* ortholog of *her6*, *deadpan* (*dpn*), is also a Notch target gene, Dpn in the *Drosophila* brain appears to promote cell cycle entry in the context of dedifferentiation (Veen, 2022).

### Prox1 in regeneration

Even though the function of Prox1 is well established within INL progenitors, here, for the first time, we demonstrate a role for Prox1 in PR differentiation. We found that Prox1 was expressed in and promoted differentiation of progenitors following PR loss. We suggest that this may occur through Liquid-liquid phase separation (LLPS). We assessed LLPS through the treatment of 1,6-hexanediol (1,6-HD). This compound dissolves phase-separates and is commonly used to determine LLPS, including a study involving the *Drosophila* ortholog of *prox1*, *prospero* (*pros*) (Liu et al., 2020). While we did observe significantly less Prox1 expression in the outer nuclear layer (ONL) following 1,6-HD treatment compared to control, to confirm LLPS is the responsible mechanism, other assays such as temperature increases could be used destabilise phase-separates (Cinar et al., 2019, Zhu et al., 2022).

### Nerfin-1/Insm1a and its target genes in regeneration

Insulinoma-associated 1a (Insm1a) has a dual function in zebrafish retinal regeneration, playing roles in both multipotency and cell-cycle exit. Upon activation via Achaete-scute complex like 1a (Ascl1a), Insm1a can function in a positive feedback-loop to increase Ascl1a expression, to activate multipotency and proliferation factors (Ramachandran et al., 2012). However, Insm1a is also essential for cell cycle exit by repressing BAF chromatin remodelling complex subunit (Bcl11) which in turn stimulates the expression of the cyclin-dependant-kinase (CDK) inhibitor: p57kip2 (Ramachandran et al., 2012). Interestingly, the *Drosophila* ortholog of *insm1a*, *nervous fingers-1* (*nerfin-1*), promotes and maintains differentiation in neurons (Vissers et al., 2018). Moreover, many targets of Nerfin-1 have known roles in zebrafish retinal regeneration. We showed that the downregulation of the Nerfin-1 target, SoxNeuro (SoxN), in the medulla neurons induced ectopic stem cells.

The zebrafish ortholog of *SoxN*, *sry* (*sex determining region Y*)-*box 2* (*sox2*), has a well-established role during regeneration. Sox2 is a multipotency factor required for the reprogramming of activated MG and its loss during regeneration decreases the proliferation of MG-derived progenitors (Gorsuch et al., 2017). Interestingly, this occurs through the activation of Ascl1a, whose *Drosophila* ortholog, *asense* (*ase*), is also a target of Nerfin-1. Together, this highlights the similarities between the genes and networks involved in dedifferentiation and regeneration as well as their conservation across model species. These similarities can be exploited to identify new factors involved in vertebrate regeneration with the potential to increase regeneration in humans.

---

**Supplementary 1.**
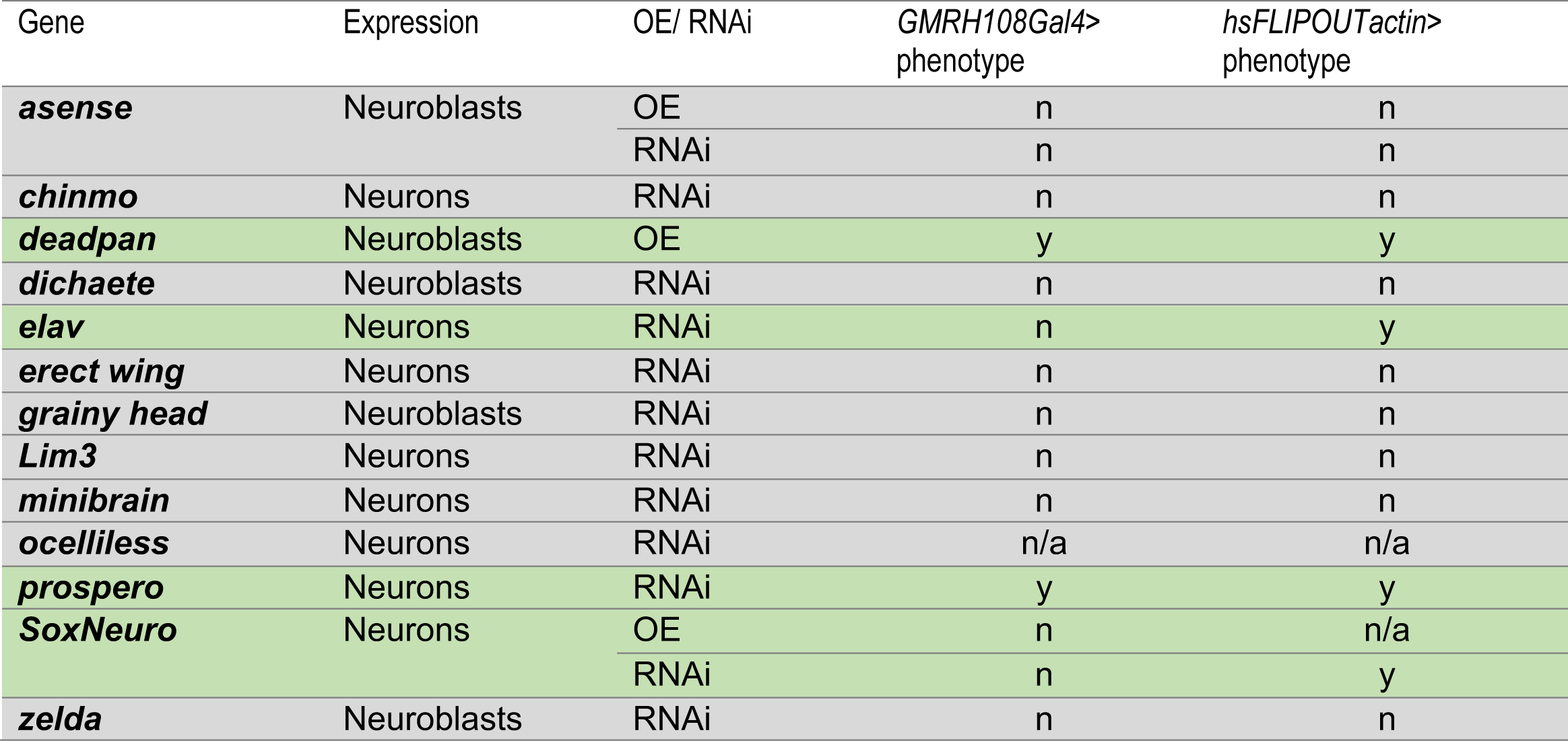
Genetic screen for candidates involved in dedifferentiation. Representative of candidates screened for involvement in medulla neuronal dedifferentiation. The name of the gene, location of expression, class of manipulation (overexpression (OE) / RNAi) and result of screen under both *GMRH108Gal4* and *hs FLIPOUTactin* promoters are shown. Grey indicates the candidate genes that were screened. White indicates candidate genes that were not screened. Green indicates that Miranda positive cells were identified. The misexpression of *deadpan* and *prospero* induced Miranda positive cells under both promoters. The knockdown of *SoxNeuro* and *elav* induced Miranda positive cells under the *hsFLIPOUTactin* promoter.

**Supplementary 2.**
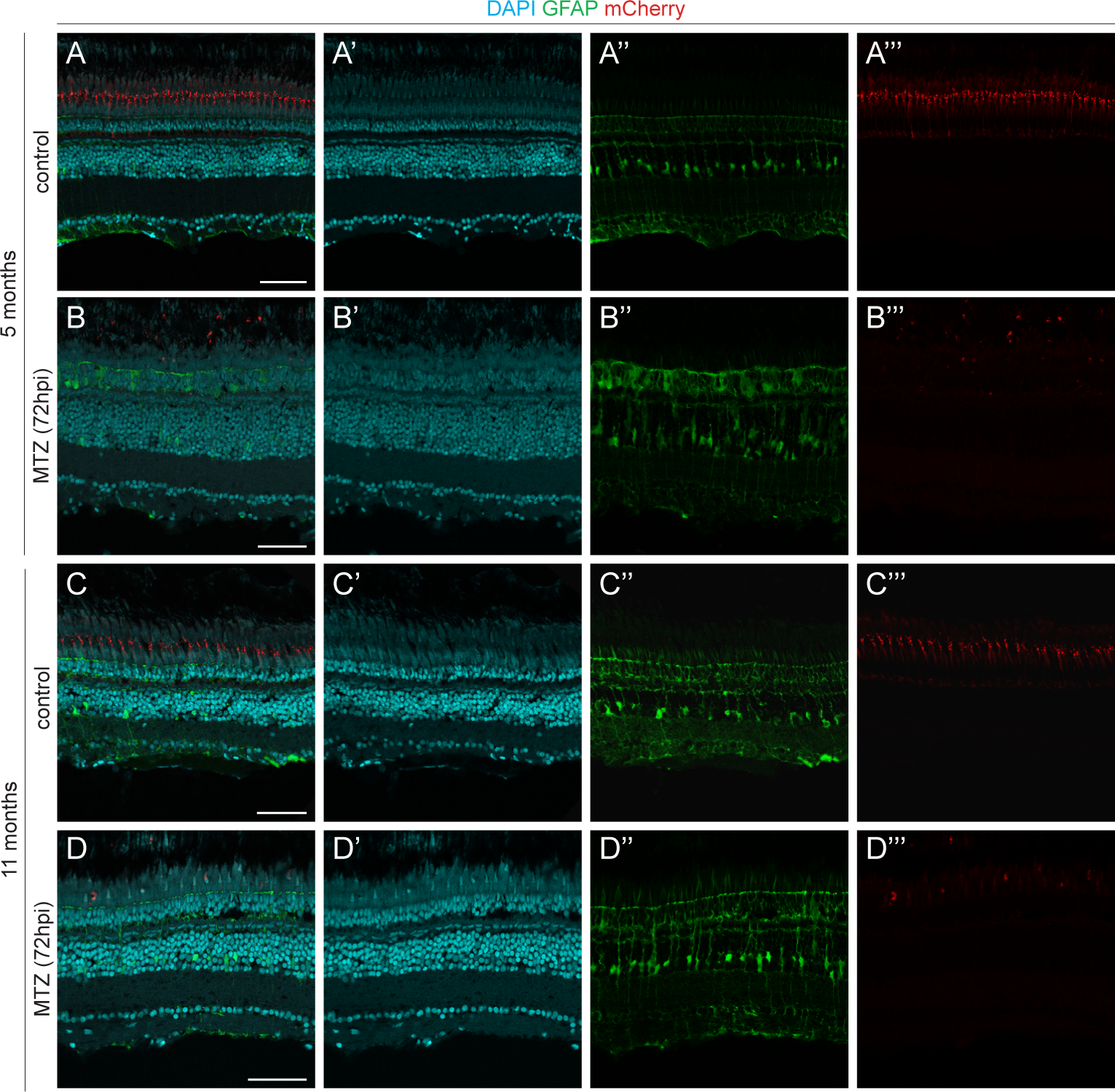
Photoreceptors are ablated using MTZ treatment in adult fish. (A-D’’’) Micrographs of retinal sections with the Inner Nuclear Layer (INL) outlined. Ablation of PRs (red) was induced with metronidazole (MTZ) treatment in Tg(*lws2:nfsb-mCherry, gfap:eGFP*) zebrafish. The control was not treated with MTZ. DAPI (cyan) labels cell nuclei, GFAP (green) labels Müller glia (MG). 5 month and 11-month-old MTZ treated fish at 72 hours post injury (hpi) had reduced mCherry compared to control. Scale bars: 50 μm.

**Supplementary 3.**
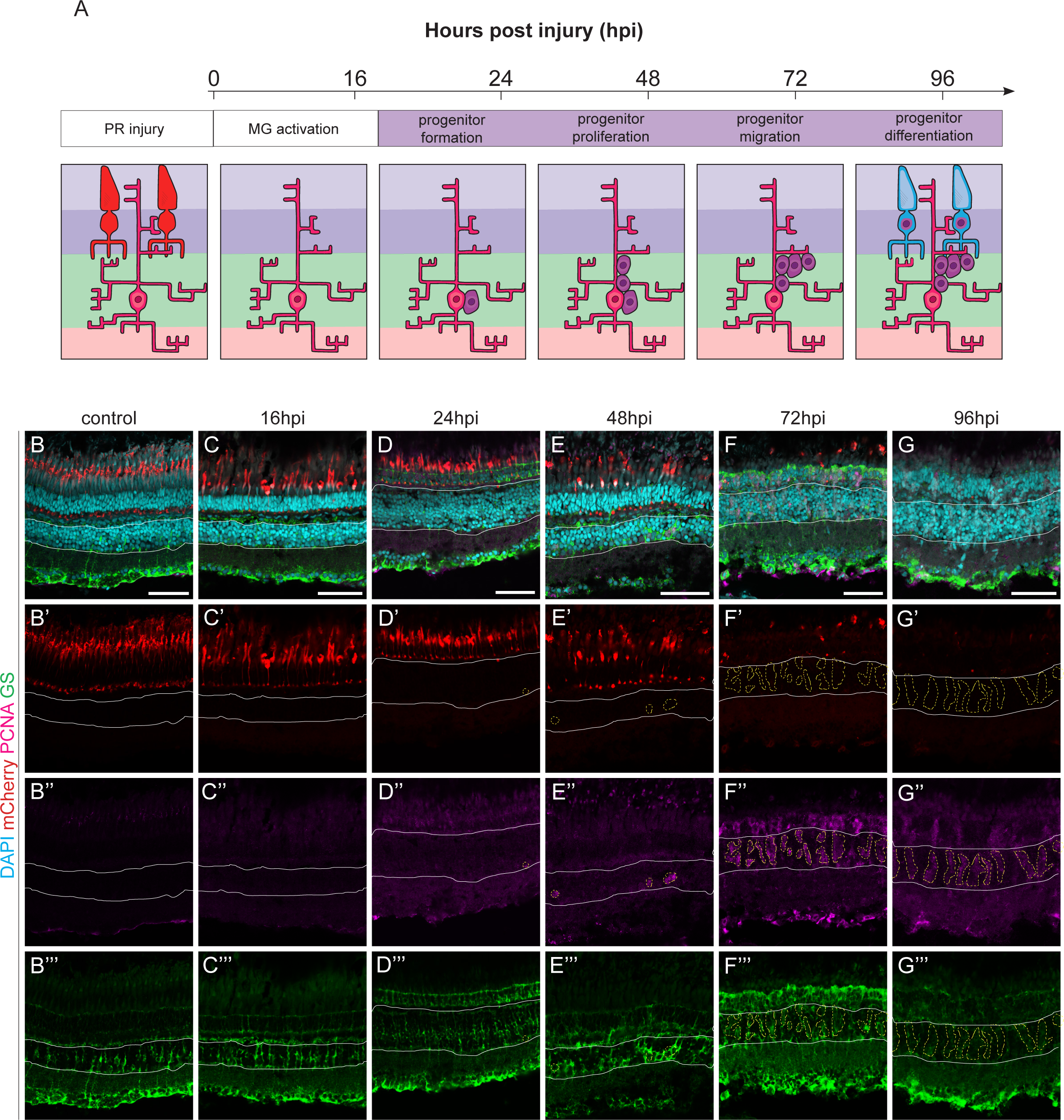
Müller glia derived progenitors drive regeneration starting 24 hours post-injury in the adult zebrafish. (A) Schematic representation of the regenerating adult retina. Time course from 0-96 hours post injury (hpi). Photoreceptors (PRs) are ablated (red, first and second panel); a Müller glia (pink, MG) derived progenitor is generated (purple, third panel); the progenitors proliferate (fourth panel) then differentiate into PRs (fifth panel). (B-G’’’) Micrographs of retinal sections with the INL outlined. Ablation of PRs (red) was induced with metronidazole (MTZ) treatment in Tg(*lws2:nfsb-mCherry*) zebrafish. Sections were stained with Proliferating Cell Nuclear Antigen (PCNA, pink) to label proliferating cells, and Glutamine Synthase (GS, green) to label MG, and DAPI to label nuclei. At 16hpi there were no PCNA+ cells in the INL (outlined), mCherry and GS expression were similar to unablated control (B-C’’’). At 24hpi and 48hpi there were single PCNA+ cells within the INL and mCherry and GS expression was similar to control (D-E’’’). By 72hpi and 96hpi there were chains of PCNA+ cells within the INL and mCherry expression was almost diminished while GS expression was diffuse compared to control (F-G’’’). Scale bars: 50 μm.

**Supplementary 4.**
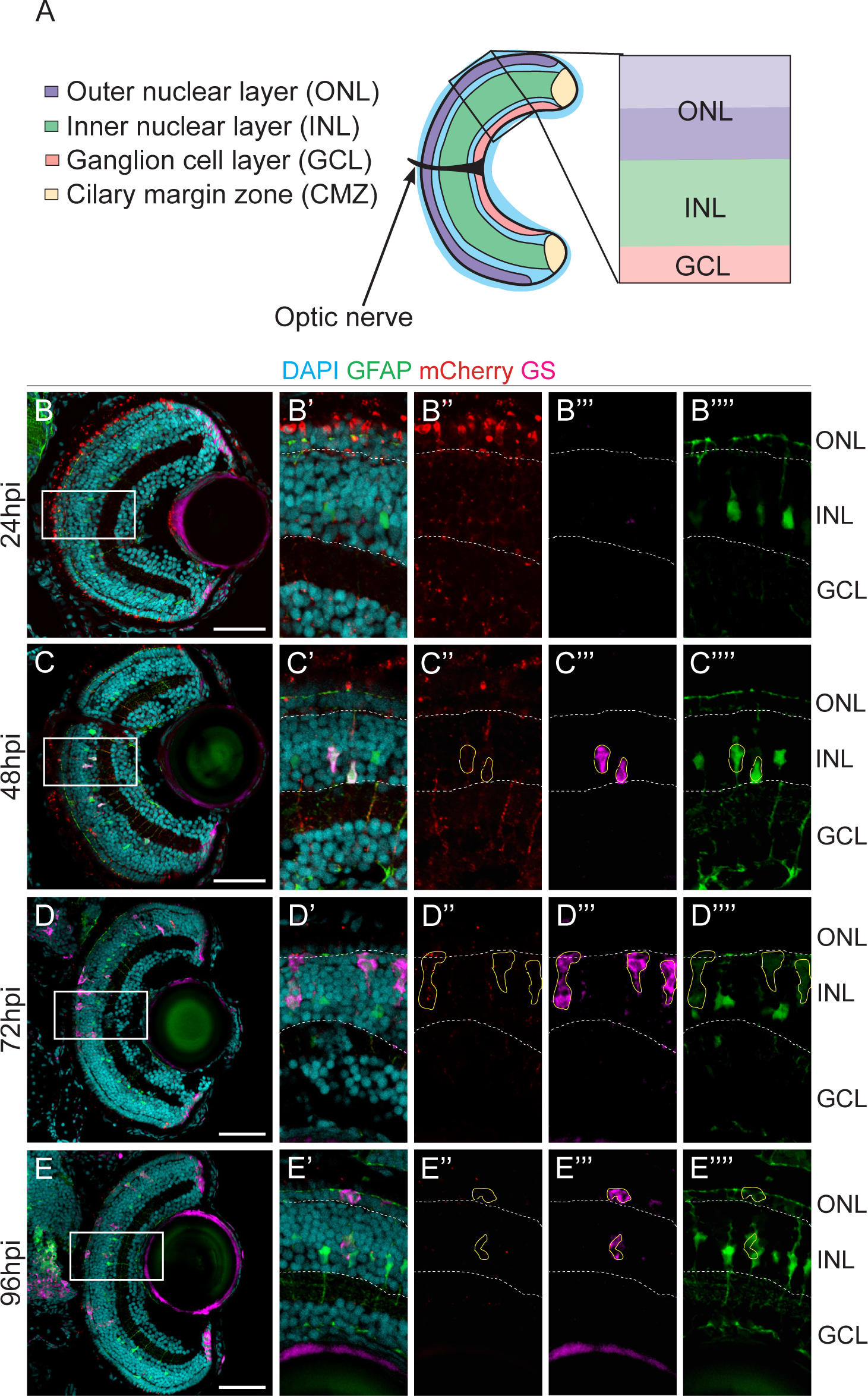
Müller glia derived progenitors drive regeneration starting 24 hours post-injury in the larval zebrafish. (A) Schematic representation of the larval retina labelling outer nuclear layer (ONL, purple), inner nuclear layer (INL, green), ganglion cell layer (GCL, pink), ciliary margin zone (CMZ, yellow) and optic nerve (arrow). (B-G’’’) Micrographs of retinal sections with the INL outlined. Ablation of PRs (red) was induced with metronidazole (MTZ) treatment in Tg(*lws2:nfsb-mCherry*) zebrafish. Sections were stained with Proliferating Cell Nuclear Antigen (PCNA, pink) to label proliferating cells, and Glutamine Synthase (GS, green) to label MG, and DAPI to label nuclei. At 16hpi there were no PCNA+ cells in the INL (outlined), mCherry and GS expression. At 24hpi and 48hpi there were single PCNA+ cells within the INL (D-E’’’). By 72hpi and 96hpi there were chains of PCNA+ cells within the INL and mCherry expression was almost diminished while GS expression was diffuse (F-G’’’). Scale bars: 50 μm.

**Supplementary 5.**
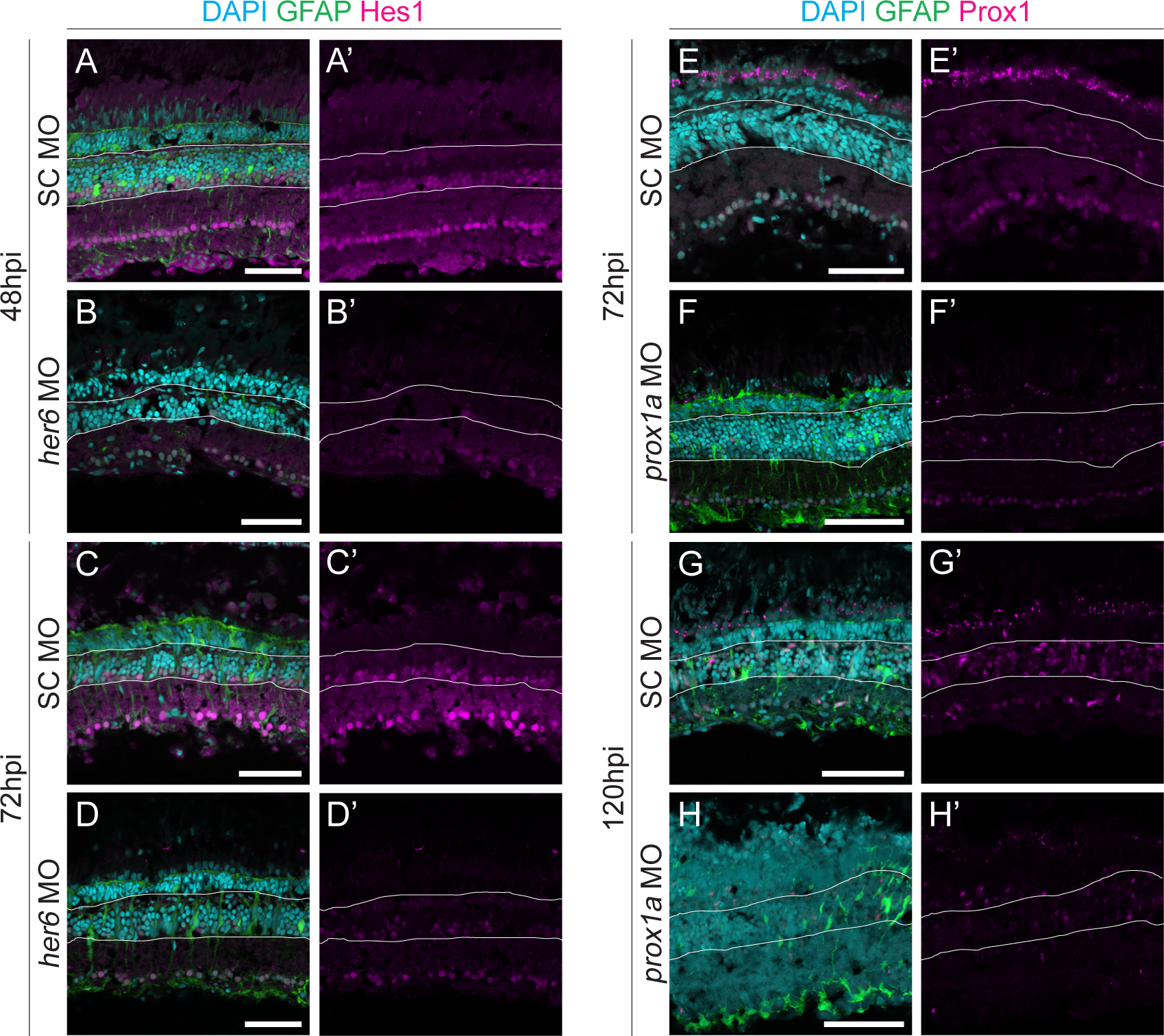
*her6* and *prox1a* MOs induce the loss of Hes1 and Prox1 protein, respectively. (A-H’) Micrographs of retinal sections comparing protein expression in SC or transgene MO injected fish. Tg(*lws2:nfsb-mCherry, gfap:eGFP*) zebrafish were injected with either a standard control (SC), *prospero homeobox 1a* (*prox1a*) morpholino (MO), or *hairy-related 6* (*her6*) MO prior to ablation. Sections were stained with Prox1 (pink) or Hes1 (pink), and DAPI to label nuclei. GFAP labels the Müller glia (green). Compared to SC MO, *her6* MO treated fish had less Hes1 expression at 48hpi and 72hpi (A-D’) compared to SC MO. *prox1a* MO treated fish had less Prox1 expression at 72hpi and 120hpi compared to SC MO (E-H’). Scale bars: 50 μm.

## References

Bernardos, R. L., Barthel, L. K., Meyers, J. R. & Raymond, P. A. 2007. Late-stage neuronal progenitors in the retina are radial Muller glia that function as retinal stem cells. J Neurosci, 27, 7028–40.

Betschinger, J., Mechtler, K. & Knoblich, J. A. 2006. Asymmetric segregation of the tumor suppressor brat regulates self-renewal in Drosophila neural stem cells. Cell, 124, 1241–53.

Campbell, L. J., Levendusky, J. L., Steines, S. A. & Hyde, D. R. 2022. Retinal regeneration requires dynamic Notch signaling. Neural Regen Res, 17, 1199–1209.

Carney, T. D., Struck, A. J. & Doe, C. Q. 2013. midlife crisis encodes a conserved zinc-finger protein required to maintain neuronal differentiation in Drosophila. Development, 140, 4155–64.

Cinar, H., Fetahaj, Z., Cinar, S., Vernon, R. M., Chan, H. S. & Winter, R. H. A. 2019. Temperature, Hydrostatic Pressure, and Osmolyte Effects on Liquid-Liquid Phase Separation in Protein Condensates: Physical Chemistry and Biological Implications. Chemistry, 25, 13049–13069.

Conner, C., Ackerman, K. M., Lahne, M., Hobgood, J. S. & Hyde, D. R. 2014. Repressing notch signaling and expressing TNFalpha are sufficient to mimic retinal regeneration by inducing Muller glial proliferation to generate committed progenitor cells. J Neurosci, 34, 14403–19.

D’orazi, F. D., Suzuki, S. C., Darling, N., Wong, R. O. & Yoshimatsu, T. 2020. Conditional and biased regeneration of cone photoreceptor types in the zebrafish retina. J Comp Neurol, 528, 2816–2830.

Duster, R., Kaltheuner, I. H., Schmitz, M. & Geyer, M. 2021. 1,6-Hexanediol, commonly used to dissolve liquid-liquid phase separated condensates, directly impairs kinase and phosphatase activities. J Biol Chem, 296, 100260.

Dyer, M. A., Livesey, F. J., Cepko, C. L. & Oliver, G. 2003. Prox1 function controls progenitor cell proliferation and horizontal cell genesis in the mammalian retina. Nat Genet, 34, 53–8.

Fogerty, J., Song, P., Boyd, P., Grabinski, S. E., Hoang, T., Reich, A., Cianciolo, L. T., Blackshaw, S., Mumm, J. S., Hyde, D. R. & Perkins, B. D. 2022. Notch Inhibition Promotes Regeneration and Immunosuppression Supports Cone Survival in a Zebrafish Model of Inherited Retinal Dystrophy. J Neurosci, 42, 5144–5158.

Fraser, B., Duval, M. G., Wang, H. & Allison, W. T. 2013. Regeneration of cone photoreceptors when cell ablation is primarily restricted to a particular cone subtype. PLoS One, 8, e55410.

Friedmann-Morvinski, D., Bushong, E. A., Ke, E., Soda, Y., Marumoto, T., Singer, O., Ellisman, M. H. & Verma, I. M. 2012. Dedifferentiation of neurons and astrocytes by oncogenes can induce gliomas in mice. Science, 338, 1080–4.

Froldi, F., Szuperak, M., Weng, C. F., Shi, W., Papenfuss, A. T. & Cheng, L. Y. 2015. The transcription factor Nerfin-1 prevents reversion of neurons into neural stem cells. Genes Dev, 29, 129–43.

Gorsuch, R. A., Lahne, M., Yarka, C. E., Petravick, M. E., Li, J. & Hyde, D. R. 2017. Sox2 regulates Muller glia reprogramming and proliferation in the regenerating zebrafish retina via Lin28 and Ascl1a. Exp Eye Res, 161, 174–192.

Hagerman, G. F., Noel, N. C., Cao, S. Y., Duval, M. G., Oel, A. P. & Allison, W. T. 2016. Rapid Recovery of Visual Function Associated with Blue Cone Ablation in Zebrafish. PLoS One, 11, e0166932.

Ile, K. E., Kassen, S., Cao, C., Vihtehlic, T., Shah, S. D., Mousley, C. J., Alb, J. G., JR., Huijbregts, R. P., Stearns, G. W., Brockerhoff, S. E., Hyde, D. R. & Bankaitis, V. A. 2010. Zebrafish class 1 phosphatidylinositol transfer proteins: PITPbeta and double cone cell outer segment integrity in retina. Traffic, 11, 1151–67.

Jorstad, N. L., Wilken, M. S., Grimes, W. N., Wohl, S. G., Vandenbosch, L. S., Yoshimatsu, T., Wong, R. O., Rieke, F. & Reh, T. A. 2017. Stimulation of functional neuronal regeneration from Muller glia in adult mice. Nature, 548, 103–107.

Kassen, S. C., Ramanan, V., Montgomery, J. E., C, T. B., Liu, C. G., Vihtelic, T. S. & Hyde, D. R. 2007. Time course analysis of gene expression during light-induced photoreceptor cell death and regeneration in albino zebrafish. Dev Neurobiol, 67, 1009–31.

Krishnakumar, P., Riemer, S., Perera, R., Lingner, T., Goloborodko, A., Khalifa, H., Bontems, F., Kaufholz, F., El-Brolosy, M. A. & Dosch, R. 2018. Functional equivalence of germ plasm organizers. PLoS Genet, 14, e1007696.

Lahne, M., Brecker, M., Jones, S. E. & Hyde, D. R. 2020. The Regenerating Adult Zebrafish Retina Recapitulates Developmental Fate Specification Programs. Front Cell Dev Biol, 8, 617923.

Liu, X., Shen, J., Xie, L., Wei, Z., Wong, C., Li, Y., Zheng, X., Li, P. & Song, Y. 2020. Mitotic Implantation of the Transcription Factor Prospero via Phase Separation Drives Terminal Neuronal Differentiation. Dev Cell, 52, 277–293 e8.

Montgomery, J. E., Parsons, M. J. & Hyde, D. R. 2010. A novel model of retinal ablation demonstrates that the extent of rod cell death regulates the origin of the regenerated zebrafish rod photoreceptors. J Comp Neurol, 518, 800–14.

Moriyama, M., Osawa, M., Mak, S. S., Ohtsuka, T., Yamamoto, N., Han, H., Delmas, V., Kageyama, R., Beermann, F., Larue, L. & Nishikawa, S. 2006. Notch signaling via Hes1 transcription factor maintains survival of melanoblasts and melanocyte stem cells. J Cell Biol, 173, 333–9.

Ng, J., Currie, P. D. & Jusuf, P. R. 2013. The Regenerative Potential of the Vertebrate Retina - Lesons from the Zebrafish. In: Pebay, A. (ed.) Regenerative Biology of the Eye. 1 ed. New York: Humana Press.

Noel, N. C. L., Macdonald, I. M. & Allison, W. T. 2021. Zebrafish Models of Photoreceptor Dysfunction and Degeneration. Biomolecules, 11.

Parmeggiani, F. 2011. Clinics, epidemiology and genetics of retinitis pigmentosa. Curr Genomics, 12, 236–7.

Pasini, A., Jiang, Y. J. & Wilkinson, D. G. 2004. Two zebrafish Notch-dependent hairy/Enhancer-of-split-related genes, her6 and her4, are required to maintain the coordination of cyclic gene expression in the presomitic mesoderm. Development, 131, 1529–41.

Pollak, J., Wilken, M. S., Ueki, Y., Cox, K. E., Sullivan, J. M., Taylor, R. J., Levine, E. M. & Reh, T. A. 2013. ASCL1 reprograms mouse Muller glia into neurogenic retinal progenitors. Development, 140, 2619–31.

Ramachandran, R., Zhao, X. F. & Goldman, D. 2012. Insm1a-mediated gene repression is essential for the formation and differentiation of Muller glia-derived progenitors in the injured retina. Nat Cell Biol, 14, 1013–23.

Raymond, P. A., Barthel, L. K., Bernardos, R. L. & Perkowski, J. J. 2006. Molecular characterization of retinal stem cells and their niches in adult zebrafish. BMC Dev Biol, 6, 36.

Soto, X., Biga, V., Kursawe, J., Lea, R., Doostdar, P., Thomas, R. & Papalopulu, N. 2020. Dynamic properties of noise and Her6 levels are optimized by miR-9, allowing the decoding of the Her6 oscillator. EMBO J, 39, e103558.

Southall, T. D. & Brand, A. H. 2009. Neural stem cell transcriptional networks highlight genes essential for nervous system development. EMBO J, 28, 3799–807.

Southall, T. D., Davidson, C. M., Miller, C., Carr, A. & Brand, A. H. 2014. Dedifferentiation of neurons precedes tumor formation in Lola mutants. Dev Cell, 28, 685–96.

Thummel, R., Bailey, T. J. & Hyde, D. R. 2011. In vivo electroporation of morpholinos into the adult zebrafish retina. J Vis Exp, e3603.

Todd, L., Hooper, M. J., Haugan, A. K., Finkbeiner, C., Jorstad, N., Radulovich, N., Wong, C. K., Donaldson, P. C., Jenkins, W., Chen, Q., Rieke, F. & Reh, T. A. 2021. Efficient stimulation of retinal regeneration from Muller glia in adult mice using combinations of proneural bHLH transcription factors. Cell Rep, 37, 109857.

Veen, F., Dong, Alvarez-Ochoa, Nguyen, Harvey, Mcmullen, Marshall, Jusuf, Cheng 2022. Cell cycle and temporal transcription factors regulate proliferation and neuronal diversity of dedifferentiation-derived neural stem cells. bioRxiv.

Vissers, J. H. A., Froldi, F., Schroder, J., Papenfuss, A. T., Cheng, L. Y. & Harvey, K. F. 2018. The Scalloped and Nerfin-1 Transcription Factors Cooperate to Maintain Neuronal Cell Fate. Cell Rep, 25, 1561–1576 e7.

Zhang, P., Yang, Y., Nolo, R., Zweidler-Mckay, P. A. & Hughes, D. P. 2010. Regulation of NOTCH signaling by reciprocal inhibition of HES1 and Deltex 1 and its role in osteosarcoma invasiveness. Oncogene, 29, 2916–26.

Zhu, S., Gu, J., Yao, J., Li, Y., Zhang, Z., Xia, W., Wang, Z., Gui, X., Li, L., Li, D., Zhang, H. & Liu, C. 2022. Liquid-liquid phase separation of RBGD2/4 is required for heat stress resistance in Arabidopsis. Dev Cell, 57, 583–597 e6.

Zhu, S., Wildonger, J., Barshow, S., Younger, S., Huang, Y. & Lee, T. 2012. The bHLH repressor Deadpan regulates the self-renewal and specification of Drosophila larval neural stem cells independently of Notch. PLoS One, 7, e46724.

